# Groove architecture controls lipid scrambling in simulations of protein and model systems

**DOI:** 10.1101/2025.06.27.662058

**Authors:** Harper E. Smith, Travis Harrison-Rawn, Wang Zheng, Angela Ballesteros, Marcos Sotomayor

## Abstract

Lipid bilayers are essential to life as they surround most cells and membrane-bound organelles. The integrity and fate of cells depend on the asymmetric makeup of lipid bilayers with various membrane proteins regulating the lipid composition of a bilayer’s two leaflets. Lipids scramblases are one of the primary regulators of lipid asymmetry in bilayers, spontaneously transferring lipids between membrane leaflets. Members of the TMEM16, OSCA/TMEM63, and TMC families have been suggested to be lipid scramblases. Despite significant differences, these proteins share a common structural architecture that features a membrane-exposed groove. The “credit card” mechanism proposes that lipids switch leaflets by moving their polar head groups either inside (partially dry) or on the surface of (wet) membrane-exposed, open hydrophilic grooves. However, emerging evidence of closed-groove scrambling challenges this model. Given the sequence diversity of groove-lining amino acids in TMEM16, OSCA/TMEM63, and TMC proteins, we hypothesized that lipid scrambling is primarily determined by groove architecture. To test this hypothesis and the credit card mechanism, we used coarse-grained molecular dynamics simulations of experimental structures and AlphaFold-generated models of six different scramblases in closed and open states. In these simulations, we observed little scramblase activity in most closed-state configurations but robust scrambling by all open-state models. We then built simplified TMEM16-based scramblases with only three bead types uniformly set for solvent-facing, transmembrane, and groove regions. We used this and further simplified models to vary groove surface hydrophilicity, groove surface geometry, and groove architecture. Our models support the partially dry and wet credit card mechanisms and suggest that groove architecture plays a more important role in facilitating lipid scrambling than the detailed sequence of groove-lining amino acids.

**STATEMENT OF SIGNIFICANCE:** Many biological processes, including blood clotting and apoptosis, are triggered by exposure of specific lipids on the outer leaflet of a lipid bilayer. Spontaneous diffusion of lipids between leaflets is slow, but scramblases can speed up transbilayer lipid exchange, a key step in signaling cascades. Here we used coarse-grained simulations to isolate and quantify the key biophysical properties of scramblases that modulate lipid movement across the bilayer. We found that the architecture of scramblases’ membrane-exposed groove is key to their function. This can be used to generate hypotheses about disease-causing mutations in known scramblases, guide drug design, and suggest favorable properties for *de novo* designed scramblases.

## INTRODUCTION

Lipid asymmetry is a fundamental property of cell membranes (1–3) that needs to be tightly regulated and that is maintained by ATP-dependent lipid translocases (4). Disruption of lipid asymmetry by scramblases is also needed and important for normal cell function and signaling (4–8). For instance, a natural asymmetry arises during lipid synthesis that requires newly produced phospholipids to rapidly move from the cytosolic to the luminal leaflet of the endoplasmic reticulum membrane (5–8). In another example, phosphatidylserine (PS), typically kept in the cytoplasmic leaflet of membranes, can be externalized to regulate blood coagulation (9) or trigger recognition of apoptotic cells (10). Spontaneous movement of lipids between leaflets happens over several hours (11), too slow for various processes that require lipid scramblases to speed up ATP-independent transbilayer lipid exchange.

The primary barrier preventing spontaneous lipid flip-flop is associated with moving the polar head group of a lipid past the nonpolar hydrocarbon tails comprising the bilayer interior. In simple models for lipid scrambling, the energy barrier (Δ*G*scramb) is reduced for smaller head groups, thinner membranes, and a higher dielectric constant in the membrane (12, 13). Lipid tails can also influence the energetics of translocation: sum-frequency vibrational spectroscopy, which can be used to track the exchange of lipids between leaflets comprised initially of either proteated or deuterated lipids (14), has been used to show that lipids with longer hydrocarbon tails have higher Δ*G*scramb for the same head group due to entropic differences. Lipids with 18C tails had around 35% higher Δ*G*scramb than lipids with 14C tails (15). The rates of lipid flip-flop also depended on membrane packing and ordering (16), which can be determined by lipid composition (17, 18) and the presence of cholesterol (19).

Lipid scramblases are proteins that disrupt the asymmetry between membrane leaflets by facilitating lipid flip-flop (scrambling) in an energy independent way (4–8, 20, 21). Scrambling occurs toward equilibrium, so existing lipid concentration gradients drive the directionality of lipid movements. A handful of scrambling protein families have been reported (22): the ATG9 proteins (23, 24) and the TMEM41B/VMP1 (25–27) families that participate in autophagy (28); the Ca^2+^-activated TMEM16 channels/scramblases (29, 30); the mitochondrial insertase MTCH2 (31) and other insertases that show scrambling activity (32); the Xk-related proteins that play a role in apoptosis (33); mitochondrial VDAC proteins (34, 35); and opsin, a GPCR in photoreceptor cells that is constitutively active with scrambling rates up to 10^5^ lipids⋅s^−1^ (36). Here we focus on the TMEM16, OSCA/TMEM63, and TMC families of scramblases and ion channels—together these proteins comprise the transmembrane channel/scramblase (TCS) superfamily, also known as TOSCA (37–56, 53, 57).

TCS proteins share a butterfly-like fold, which includes 10 or 11 transmembrane alpha helices (13, 38, 39, 43, 44, 48, 51, 54, 58–82) labeled α0 to α10. Most superfamily members have several short extracellular loops, but their cytoplasmic domains differ: TMEM16s contain N- and C-terminal cytoplasmic domains with mixed α/β topology (13, 38, 68–82); OSCA/TMEM63s feature long, membrane-parallel helices (beam-like domains) ending in membrane-embedded loops (hooks) (51, 54, 58–67); and TMCs interact intracellularly with Ca^2+^-binding CIB proteins (43, 44, 46, 83). Interestingly, structures of some TCS members reveal a hydrophilic, membrane-exposed groove formed by α3 through α7 (13, 38, 53, 59, 64, 69, 73, 78) that may enable water/ion permeation and lipid scrambling. Moreover, cryo-EM reconstructions of some TCS family members suggest that the exposed groove is lined by lipids (13, 53, 59, 78), creating a proteo-lipidic pore that facilitates ionic currents varying with membrane composition (53). Structures for other TCS family members have revealed closed grooves. Although TCS family members have differing sequences, the overall structural architecture of open and closed grooves seems to be conserved.

Scramblases are traditionally thought to lower Δ*G*scramb by accommodating hydrophilic head groups through an open, membrane-exposed hydrophilic groove while the hydrophobic tail passes through the membrane interior (84). According to the original credit card mechanism (47, 76, 84), scrambled lipids fully insert their hydrophilic head groups into the groove, forming direct protein-head group interactions (partially dry mechanism), although there is evidence that lipid headgroups may pass on the surface of a hydrated groove (wet mechanism) (76, 85–91). However, evidence conflicting with the credit card mechanisms has emerged: *Aspergillus fumigatus* (af) TMEM16 is able to scramble PEGylated PE derivatives with diameters up to 42 Å (92), and mutations in groove-lining residues in afTMEM16 minimally affect scrambling rate (13). This evidence suggests that scrambling does not occur through the groove, but rather in a nonspecific way governed by local membrane properties like thinning (59, 69, 76, 92, 93) or curvature (43, 53, 59, 73, 88, 93).

Structures and models of TCS proteins in both open and closed states are now available, paving the way for a systematic analysis of the determinants of lipid scrambling throughout the family. The groove formed by α3 through α7 is enlarged in structural models of open-like states of TMEM16s (13, 38, 69, 73, 78), OSCAs (53, 54, 61, 64), TMEM63A (57), TMEM63B (59), and TMCs (46, 50, 94). Despite sequence differences, experiments and simulations have shown that members of these families can scramble under activating conditions (13, 37, 49, 59, 69, 70, 73, 77–79, 87, 91, 94–105). Additionally, Martini3 (106) coarse-grained (CG) simulations have correctly predicted scrambling activity by TMEM16F, TMEM41B, VMP1, ATG9, VDAC1, VDAC2, rhodopsin, MCP1, and insertases (32), but no scrambling for negative controls (32); similar results were obtained for TMEM16 proteins (91). Here, we systematically examined scrambling using Martini3 models of TCS proteins. We found that all simulated putative open states were scrambling-competent and that closed states were mostly not, except for the closed state of TMC1 which scrambled lipids at moderate rates. Most scrambling events occurred with headgroups occupying the surface of the hydrated open grooves (wet credit card mechanism) and in rare cases we observed noncanonical scrambling, i.e., scrambling through a dimeric interface or other non-groove surfaces. To test the physico-chemical determinants of lipid scrambling, we used TMEM16-based simplified models with varying hydrophilicity and surface geometry of groove residues, as well as artificial cubic scramblases with varying groove architectures. Simulations of these models predict that scrambling activity is strongly controlled by groove architecture and hydrophilicity, but not fully dependent on the exact sequence of amino acids lining the groove.

## METHODS

### TCS family scramblases

We prepared all-atom scramblase models based on closed state *Nectria hematococca* (nh) TMEM16 (PDB: 6QM4 (78)), an open-like state of nhTMEM16 (PDB: 4WIS (38)), closed state hsTMEM63A (PDB: 8EHW (51)), closed state hsTMEM63B (PDB: 8EHX (51)), open-like hsTMEM63B (PDB: 8WG4 (59)), an AlphaFold2 (AF2) model of closed state hsTMEM63C (107), closed state *Arabidopsis thaliana* (at) OSCA1.2 (PDB: 6MGV (60)), open-like atOSCA1.2 (PDB: 8XAJ (53)), a single subunit of dimeric closed-state atOSCA1.2, AF2-Multimer (108) (AF2m) closed models of dimeric *Homo sapiens* (hs) TMC1, AF2m closed models of dimeric hsTMC1 with two copies of hsCIB2, and an AlphaFold3 (109) (AF3)-based open-like model of dimeric hsTMC1 from targeted molecular dynamics simulations (94) after replacing the closed-state subunit with a second copy of the open-like configuration by alignment (Figure 1, S1). For models based on cryo-EM or crystal structures, missing loops were added by MatchMaker alignment of AF2 predictions from the AlphaFold Database (AFDB) (110) in ChimeraX (111). Missing loops were placed after alignment of the residues flanking the missing region (starting at 20 flanking residues but adjusting when needed for good alignment). Proteins were oriented in the membrane using the PPM 3.0 webserver (112) or CHARMM-GUI (113) and embedded in pure POPC. TIP3P water (114) was added to create 18 Å of padding between protein and its periodic image in the *z*-direction. Systems were neutralized with 150 mM KCl. Protonation states of neutral histidine residues were chosen by inspection and disulfide bonds were added when cysteine residues were sufficiently close.

**Figure 1.**
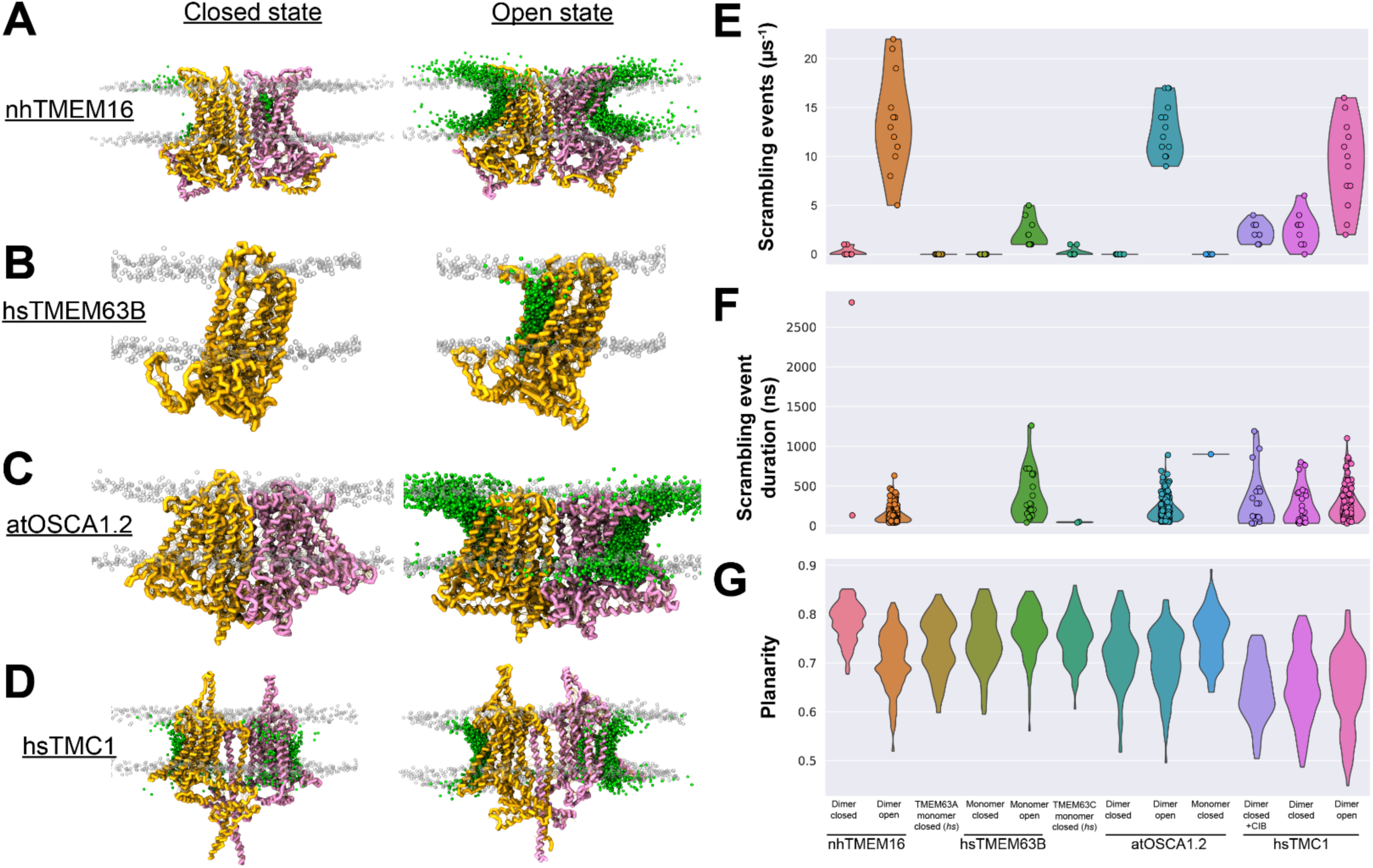
Scrambling in CG Models of Transmembrane Channel/Scramblase Superfamily Members. (*A*– *D*) Martini3 CG models of nhTMEM16 in the closed (PDB: 6QM4) and open (PDB: 4WIS) states; hsTMEM63B in closed (PDB: 8EHX) and open (PDB: 8WG4) states; atOSCA1.2 in closed (PDB: 6MGV) and open (PDB: 8XAJ) states; and AlphaFold2/AlphaFold3-based models of hsTMC1 in closed and open states. Protein models are shown in yellow and pink, non-scrambling head groups in transparent gray, and positions of head groups throughout each scrambling event are overlaid in green spheres. Models in (*A*–*D*) appear in Figure S1. (*E*–*G*) Quantification of lipid scrambling rate, duration per event, and lipid planarity; values closer to 1 are more planar.

### All-atom MD simulations

After rebuilding missing loops using AF2 predictions, we generated all-atom models for coarse graining by performing short atomistic molecular dynamics (MD) pre-equilibrations. The AF3-based open model of hsTMC1 was equilibrated as described in (94). hsTMEM63A, hsTMEM63B, and atOSCA1.2 simulations were equilibrated as described in (51). All-atom simulations were performed in NAMD 2.14 (115) using the CHARMM36 force field (116) with the CMAP correction. The SHAKE algorithm was used with fixed hydrogens and a 2 fs timestep. Electrostatic interactions were computed every other timestep using the particle mesh Ewald method. Van der Waals interactions were truncated at 12 Å using a switching function beginning at 10 Å and a Langevin piston with a damping coefficient γ = 1 ps^−1^ was applied. Systems (excluding hsTMEM63A, hsTMEM63B, atOSCA1.2, and open-like hsTMC1) were subjected to four stages of pre-equilibration: 1) 1,000 steps of minimization and 0.5 ns of MD with all atoms except lipid tails fixed; 2) 1,000 steps of minimization and 0.5 ns of MD with constraints applied to all protein backbone atoms; 3) 0.5 ns of MD with constraints applied to protein backbone atoms excluding the AF2-rebuilt loops; 4) 0.5 ps of MD with no constraints applied. To retain details from the initial model but minimize steric clash, positions were saved after stage 4 and used for coarse graining.

### Martini3 protein models

After all-atom minimization and equilibration, atomistic water, ions, and lipids were discarded and protein models were mapped to Martini3 (106) resolution using Martinize2/Vermouth (117) (vermouth-0.10.1.dev93) with secondary structure computed by DSSP in MDTraj. For TCS family systems, elastic network model bonds were added (*k* = 700 kJ⋅mol^−1^⋅nm^−2^, lower bound 5 Å, and upper bound 8 Å). For models with multiple subunits, elastic bonds were added between subunits using the same distance criteria. Additional molecules were added in Insane (118) (insane-1.2.0): a lipid bilayer was included, regular size water beads (bead W) were placed to reach dimensions around 185 × 185 × 220 Å^3^, and monovalent ions (beads NA and CL) were added to neutralize and reach a final ionic concentration of 150 mM (Supplemental Tables). The *z*-position in the membrane was specified to match placement by PPM 3.0.

### Simple protein-based scramblase models

Simple systems were constructed starting from the Martini3 representation of open state nhTMEM16 (based on PDB: 4WIS (38)). Position restraints were used for all protein beads (*k* = 5,000 kJ⋅mol^−1^⋅nm^−2^). To improve numerical stability in simulations, all bonds, angles, and dihedrals involving only protein beads were removed and exclusions were added between all pairs of protein beads within 15 Å; this effectively removes all intra-protein interactions. The virtual atom in each tryptophan (bead name SC3) was given a mass of 36 amu so their interactions could be excluded. Bead types and charges were modified using tools from the Martinize2/Vermouth library (117) in a way that retains bead size (tiny, small, or regular). For protein beads changed to Q-type, charges were set to 0 to maintain a reasonable net protein charge and prevent an excess of cations/anions during system charge neutralization. Simple models with unique non-groove transmembrane (TMng) and/or groove regions were prepared using the MDAnalysis and Biopython selection languages in combination with Martinize2/Vermouth ITP writers. TMng regions were defined using boundaries in the *z*-direction calibrated to match membrane placement in PPM 3.0. For models with uniform beads in the groove region, forty-eight groove-lining beads were selected by inspection of the protein surface as those along the groove between TM helices α3–α7 of the open-state nhTMEM16.

### Generating alternative pockets

Hypothetical membrane-spanning pockets with unique geometry and similar surface area to the original groove were generated as follows. The 48 beads in the original pocket were rotated in the *xy*-plane about the geometric center of the nhTMEM16 dimer by 20 to 160° to give new virtual groove positions. The rotated groove positions sometimes lie inside the protein, so they were translated 50 Å away from the center of the protein. The transformed positions were projected onto the protein’s surface using the Hungarian algorithm in SciPy, which found optimal pairings to minimize distances between the transformed positions and protein beads. From this new surface patch, a graph with a distance-based adjacency list (cutoff 5 Å) was constructed in NetworkX. This sometimes resulted in a spatially disconnected patch of residues, so the largest connected component was retained. To assign up to 48 beads to the new groove-like patch, beads were sequentially added vertically using an expanding search radius until the membrane region was spanned. If 48 beads were still not assigned, we fit a plane to the beads selected so far, then iterated over the lowest-degree nodes in the growing groove-like patch (those near the patch boundary) and added the neighbors closest to the plane.

### Cubic scramblases

A cube of side length 80 Å was constructed from regular size Martini3 beads (diameter 4.7 Å) using MDAnalysis. As with the simple protein models, exclusions were generated for all pairs within 15 Å and absolute position restraints were added (*k* = 5,000 kJ⋅mol^−1^⋅nm^−2^) using routines from Martinize2/Vermouth (117). The thickness of the TM region was calibrated to match the simple TMEM16-based systems, and the groove thickness in the *xy*-plane was 15 Å (3 beads). Groove and solvent-facing regions were comprised of bead type P2 while hydrophobic TMng beads were C4. Four identical grooves were placed at the center of each cubic face in the TM plane by removing beads. Spherical, ellipsoidal, cuboidal, cylindrical, and ramped grooves were constructed according to basic geometric definitions. Each shape has a corresponding shape factor *s*; larger values of *s* lead to larger pores. The cubic scramblases were subjected to the same solvation, ionization, and membrane building protocol in Insane (118) used for Martini3 protein models.

### CG simulations

CG MD simulations were performed in GROMACS (119) 2024.2 using the *New-RF* parameter set (120), including a 20 fs timestep and reaction field (RF) electrostatics. Pressure was maintained at 1 bar (compressibility 3 ⋅ 10^−4^ bar^−1^) semi-isotropically with the Berendsen barostat (121) (coupling constant 4 ps^−1^) and temperature was held at 310 K by the v-rescale thermostat. To avoid artifacts arising from too-infrequent neighbor list updates (122), we disabled *verlet-buffer-tolerance*, adopted a conservative neighbor list cutoff of 1.5 nm, and set *nstX* variables to 20 steps (as recommended for *New-RF*). Relaxation of the system was achieved by following the Martinate protocol (123): we sequentially performed up to 500 steps of position-restrained energy minimization in vacuum; up to 500 steps of position-restrained energy minimization after water, ions, and lipids had been added; 10 ps of position-restrained *NVT* with a 2 fs timestep; and 25, 50, and 300 ps of unrestrained *NpT* with timesteps of 5, 10, and 20 fs, respectively. Simulations were extended to at least 10 µs (Supplemental Tables) and positions were recorded every 100 ps. Simulations (over 150) totaled over 1.5 milliseconds.

### Quantification of lipid scrambling

Lipid scrambling was computed based on an established procedure (32, 93). Before analysis, protein atom positions were unwrapped and centered (without alignment), then non-protein atom positions were wrapped back into the periodic box. We then removed rotation of the protein about the *z*-axis. For all lipids, a vector was drawn from the end of each tail to the head group (for POPC, bead C4A/C4B to NC3), and the angle of those vectors with respect to the *z*-axis was recorded. Using this definition, lipids in the cytoplasmic and exoplasmic leaflet should approach 0° and 180°, respectively. Lipids with an average orientation angle less than or greater than 90° were initially assigned to the cytoplasmic or exoplasmic leaflet, and the running average of the lipid orientation angle was computed over 100 ns. Scrambling events were counted when a lipid from the cytoplasmic (exoplasmic) leaflet crossed 145° (35°), after which the lipid was considered to belong to a new leaflet. Once a scrambling event was identified, we determined a more accurate transition time using a 10-ns running average of the angle and more conservative cutoffs of 27.5° and 152.5°. For plotting, we divided the simulations into 1-µs segments and reported how many scrambling events were completed within each segment. Scrambling events are reported as net values for entire systems when applicable and unless otherwise indicated. Over 5,000 events were monitored throughout all simulations. Timelapse images were rendered in VMD using the built-in trajectory display setting for representations with lipids shown every 10 ns.

### Quantification of planarity

We used a two-step procedure to find the two planes that define a bilayer when lipids do not partition cleanly into leaflets. First, the RANSAC regressor from Scikit-Learn with a linear regressor was used to fit a plane to all lipid head group positions. This algorithm reduces the distance between the plane and head groups in one of the leaflets, which defined the first plane. All head groups within 5 Å of this first plane were removed from consideration, then a second plane was fit to the remaining points. The fraction of head groups falling within 5 Å of either plane was defined as the planarity, which was accumulated over the final 8 µs of simulation for plotting. Well-ordered bilayers exhibit planarity values near one.

### Quantification of averaged membrane *z*-positions

We partitioned bilayers into two leaflets using the LeafletFinder tool in MDAnalysis, then recorded head group *z*-positions (interval 10 ns) from the centered trajectories for members of each leaflet using the MembraneCurvature module in MDAnalysis. Head groups of lipids that participated in complete scrambling events were excluded from both leaflets. Based on alignment of protein beads only, rotations of the protein about the *z*-axis were removed. We binned head group *z*-positions into 100 × 100 grids, then applied a circular mask with the same diameter as the minimal *xy*-extent of the box to exclude artifacts due to alignment. Plots were constructed in Matplotlib with Gaussian interpolation applied. Grid points that were empty during any frame appear gray. We report membrane thickness as the distance between upper and lower leaflet head group *z*-position heatmaps, which were defined on the same grid. If a bin had an undefined *z*-position for either leaflet, the corresponding thickness was also undefined and appears gray in our plots.

### Water and lipid densities

Densities of phosphate atoms, positively and negatively charged ions, and water molecules in CG simulations were computed using VMD’s VolMap tool using Martini 3 radii (set via the radscale option; 1.56 Å for both water and name PO4 and 1.46 Å for ions). Volmap was used in density mode, which applies a Gaussian smoothing to the mass-weighted atomic coordinates. Densities were computed using frames every 10 ns and visualized in VMD using density thresholds of 0.033 (head groups), 0.058 (water), and 0.001 or 0.01 Å^−1^ (ions). Densities were read using the GridDataFormats package, then densities were considered to be overlapping in a voxel if both densities exceeded the threshold in that voxel.

### Statistical analyses

Unless otherwise noted, all reported scrambling rates, durations per event, and planarity values are mean ± margin of error, where the margin of error corresponds to the 95% confidence interval (CI).

## RESULTS

### Scrambling by CG models of TCS family proteins

To gain a molecular view of how TCS proteins scramble lipids, we collected structures from the protein data bank (124) or AFDB (110) and systematically converted to CG resolution. The Martini3 CG mapping procedure replaces each amino acid with ∼3-4 beads on average, based on the chemical properties of each residue, speeding up MD simulations by reducing the number of particles compared to all-atom models. We generated CG models of nhTMEM16, atOSCA1.2, hsTMEM63A–C, and hsTMC1 with or without hsCIB2, a cytoplasmic binding partner of hsTMC1 (46, 83) (Figure 1A–D, S1). After embedding each protein in a pure POPC bilayer, we simulated each system for at least 10 μs (Table S1) and monitored lipid scrambling events (Movies S1–S9). Scrambling events were identified by tracking the orientation angle of each lipid with respect to the *z*-axis over time. To visualize scrambling pathways, we extracted the positions of lipid head groups (name PO4) in the process of scrambling, taking positions every 10 ns, and overlaid them on the corresponding protein structures (Figure 1A–D, green).

Determining whether models of TCS proteins represent scrambling-competent states, states permeable to ions and water only, or semi-open states is difficult. To validate our methodology, we compared the first known open state structure of a TCS protein, nhTMEM16 (38), to a corresponding closed state (78) (Figure 1A). We recorded 13.6 ± 3.7 lipids⋅μs^−1^ (95% CI) for open state nhTMEM16 and little scrambling for its closed state (0.2 ± 0.3 lipids⋅μs^−1^, Figure 1A, 1E). We observed lipids passing through the groove between α4 and α6 on both protomers (Figure 1A, S1, Movie S1), in agreement with previous Martini3 simulations of TMEM16s in DOPC (91), Martini2 simulations of nhTMEM16 (125) and hsTMEM16K (79), and all-atom simulations of nhTMEM16 (85, 88, 89, 93) and mmTMEM16F (86, 87, 126). We estimated the time per scrambling event by tracking how long the average lipid orientation angles remained between 35 and 145°; this revealed that scrambling events for the nhTMEM16 open state occur over 134 ± 15.9 ns (Figure 1F). Most of the scrambling events involved lipids passing through the open and hydrated nhTMEM16 groove (Figure 2A, Movie S1), with some rare events involving noncanonical lipid flip-flop at the dimer interface (Figure 2A, Movie S2). These results confirmed that our models can predict lipid scrambling activity in known scramblases.

**Figure 2.**
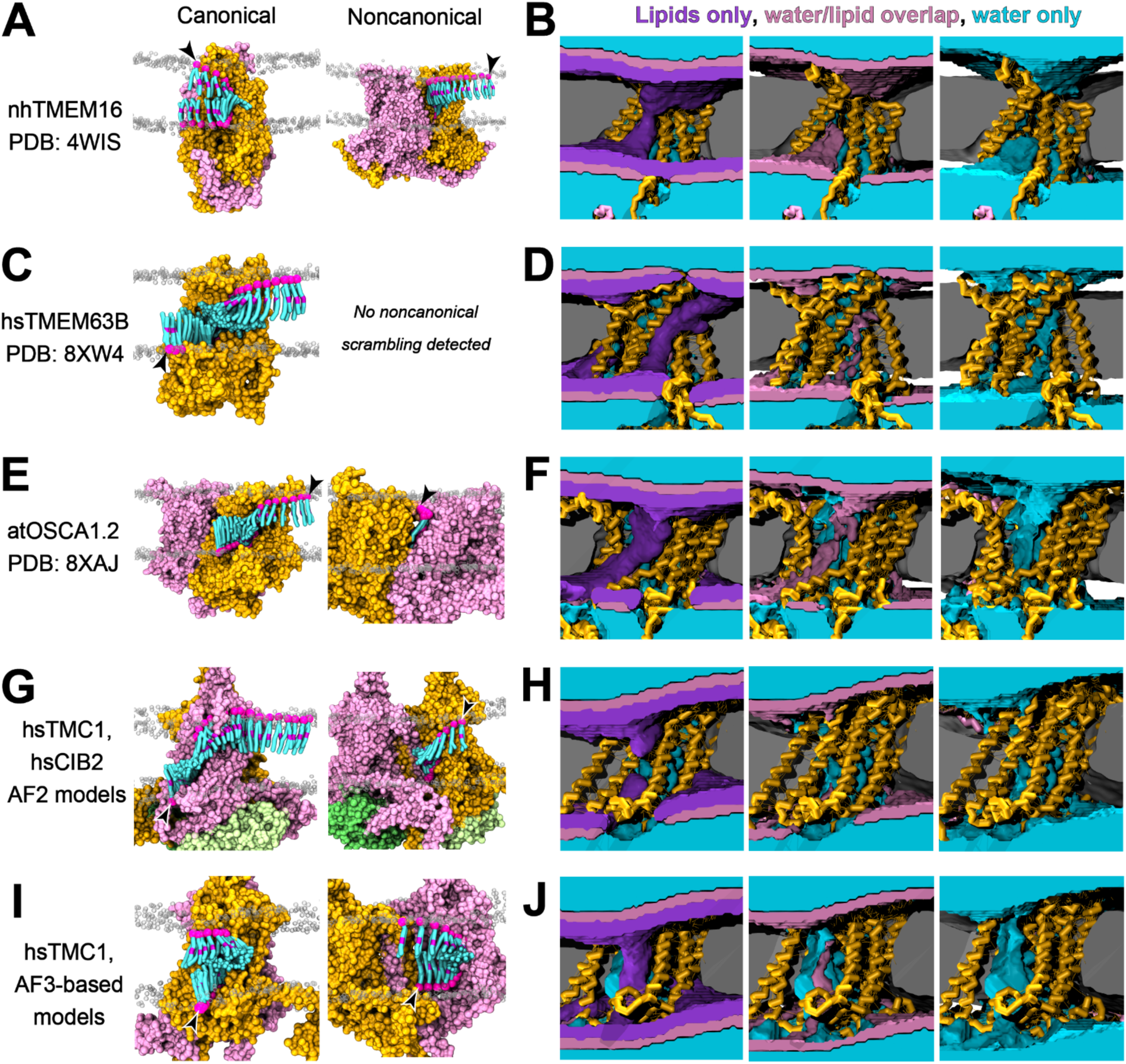
Scrambling pathways with water and head group densities. *(A)* Canonical (*left*) and noncanonical (*right*) scrambling events observed for dimeric open-state nhTMEM16 (PDB: 4WIS). Protein subunits are shown in pink and orange surface, scrambling lipids are cyan and pink, and non-scrambling head groups are transparent gray. In timelapse renders, lipids are shown every 10 ns. (*B*) Water (*left*; cyan) and lipid head group (*right*; purple) densities are shown alongside regions where the densities overlap (*middle*; pink). (*C*–*H*) Analogously, we show monomeric open-state hsTMEM63B (PDB: 8WG4) (*C, D*), dimeric open-state atOSCA1.2 (PDB: 8XAJ) (E, F), dimeric closed hsTMC1 with hsCIB2 (AF2 model) (*G*, *H*), and dimeric open hsTMC1 (AF3 model) (*I, J*). Arrowheads mark the position of the lipid at the beginning of the scrambling event. Timelapse images correspond to Movies S1-S9.

Another member of the TCS family that has been shown to scramble lipids under activating conditions is hsTMEM63B (59). In contrast to TMEM16, OSCA, and TMC family members, TMEM63 proteins function as monomers (51, 58, 62). An open state model of hsTMEM63B (59) scrambled 2.1 ± 1.1 lipids⋅μs^−1^ through the groove formed by α3–α7 (Figure 2C, Movie S3), but we observed no scrambling for the closed state of hsTMEM63B (Figure 1B, 1E). In separate work with open-like structural models of hsTME63A, we monitored scrambling at 1.9 ± 1.4 lipids⋅μs^−1^ for the wildtype protein and 5.8 ± 1.4 for a gain-of-function mutant, p.V53M (57). Experimental scrambling rates for hsTMEM63s have not been reported, but a low scrambling rate for the open state of hsTMEM63B compared to nhTMEM16 is consistent with the smaller open-state groove in the cryo-EM structure of hsTMEM63B (38, 59) (Figure S2). In this case, all scrambling events involved lipids passing through the open and hydrated hsTMEM63B groove (Figure 2D).

Next, we turned to OSCAs, plant relatives of TMEM63s that form dimers (60, 63–67). For atOSCA1.2, which has a larger groove than hsTMEM63B in open-like cryo-EM structures (53, 59) (Figure S2), we detected scrambling at 12.6 ± 1.8 lipids⋅μs^−1^ in the open state but no scrambling in the closed state (Figure 1C, 1E). As with hsTMEM63B, lipids passed through the hydrated open groove (Figure 2E, 2F; Movie S4) with some lipid translocation away from the canonical groove (Figure 2E, 2F; Movie S5). Scrambling has not been reported in wildtype OSCAs, but they share high structural similarity with TMEM63s/TMEM16s and can be converted into constitutively active scramblases (37). These simulations suggest that open state models of TCS family members can scramble lipids because they share a conserved groove architecture.

Consistent with their classification as closed states, we monitored little to no scrambling for the closed states of nhTMEM16, hsTMEM63A, hsTMEM63B, hsTMEM63C, and atOSCA1.2 (Figure 1A–E, S1). The exception was for AF2 models of dimeric closed state hsTMC1, for which we calculated 2.5 ± 1.4 lipids⋅μs^−1^ and 2.2 ± 0.7 lipids⋅μs^−1^ in the absence or the presence of hsCIB2, respectively (Figure 1D, 1E). These results suggest that hsCIB2 does not regulate closed-state scrambling activity of hsTMC1. Most scrambling occurred through the closed canonical groove for hsTMC1 in the presence of hsCIB2 (Figure 2G, 2H, Movie S6), but we detected some rare events near the dimeric interface (Figure 1D, 2G, 2H, Movie S7). The AF3-based open-like model of hsTMC1 scrambled 8.9 ± 3.0 lipids⋅μs^−1^ (Figure 2I, 2J, Movies S8 and S9), suggesting that our closed state may represent a semi-open configuration, that hsTMC1 is a constitutively active scramblase that can be further activated into a fully scrambling state, or that additional factors regulate lipid scrambling by TMCs.

It has been proposed that lipid scrambling can be catalyzed by membrane deformation (13, 43, 59, 69, 73, 76, 88, 91–93), which could decrease the distance traversed by the polar head groups through hydrophobic regions during scrambling. We defined the “planarity” of lipids throughout each simulation as the fraction of head groups within 5 Å of two best-fitting planes, which tests how well head groups follow a bilayer-like distribution throughout the final eight microseconds of simulation time (Figure 1G). Surprisingly, open and closed states of the same protein tended to exhibit similar planarity distributions, suggesting that planarity alone is a poor predictor of scrambling rate (Figure 1G). An exception is nhTMEM16, which induced a decrease in planarity in the scrambling-competent state (planarity 0.70 ± 0.07; mean ± SD) with respect to the closed state (planarity 0.78 ± 0.04; mean ± SD) (Figure 1G). Local curvature, which can be observed in the heatmaps of per-leaflet lipid head group *z*-positions, revealed membrane bending both near and far from each protein (Figure S3A–L); one notable case is nhTMEM16, which appears to more strongly deform both leaflets in the open state (Figure S3A, S3B). In contrast, hsTMEM63B and atOSCA3.1 induce similar local curvature in scrambling competent and incompetent states (Figure S3D, S3E, S3G, S3H). We observe the most pronounced distortions for TMCs (both with and without CIB2) (Figure S3J–L). Membrane thickness, which we define as the distance between head group *z*-position heatmaps between the upper and lower leaflet, decreased in proximity to the protein but was uniform far from the protein in most cases (Figure S3A–L), regardless of scrambling activity. In summary, our simulations did not demonstrate a strong correlation between scrambling activity and either membrane thickness far from the protein or curvature. Actively scrambling proteins exhibited decreased membrane thickness in the immediate vicinity of the groove, consistent with results for afTMEM16 (13).

### Constructing simple CG scramblases with uniform beads

Our CG simulations of TCS family members revealed that lipid scrambling can be facilitated by proteins that share a similar open-state groove architecture, independent of sequence variation. However, the scrambling rates observed for our Martini3 models of TCS family proteins are likely affected by various biophysical properties besides groove architecture, including polarity and geometry of surface residues in the groove. To isolate and evaluate the impact of these factors, we built simple scramblases with the same groove architecture as the open state of nhTMEM16 (38) (Figure 3A–C). Unlike our previous models, these simple scramblases have positional restraints to retain their initial shape (see Methods). Without changing overall architecture or surface geometry, we then sought to replace the complex pattern of surface hydrophilicity in nhTMEM16 (Figure 3C) with a simple set of Martini3 “beads” (CG particles) (Figure 3D) that could maintain scrambling activity.

**Figure 3.**
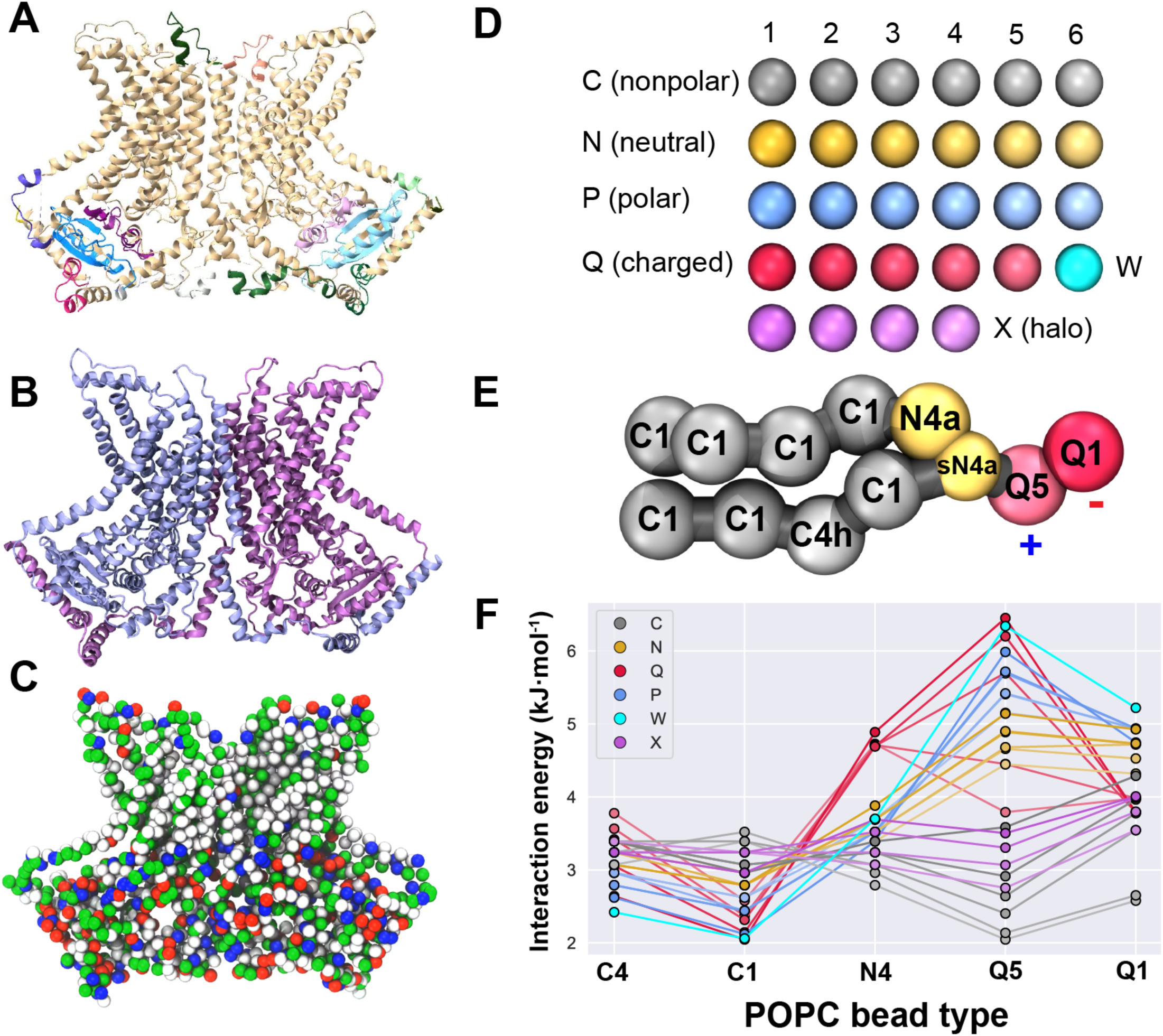
Constructing simple model scramblases based on nhTMEM16. (*A*) Cartoon representation of nhTMEM16 (PDB: 4WIS; tan) with loops from an AlphaFold2 model in colors. (*B*) nhTMEM16 after a brief simulation. Subunits of the dimer are blue and purple. (*C*) CG representation of nhTMEM16 with beads colored by residue type: hydrophobic beads are white, polar are green, positively charged are blue, and negatively charged are red. Only backbone beads are shown. (*D*) Schematic representation of the bead types available in Martini3. (*E*) CG POPC with apolar tails (C-type), intermediate/nonpolar glycerol backbone (N-type), and charged head group (Q-type). Glycerol beads have hydrogen bond acceptor labels as suffixes (a) and the sN4a bead is small size (s prefix). In bead type C4h, the “*h*” suffix is a label that increases self-interaction. (*F*) Martini3 interaction levels between POPC beads and all other bead types. Bead subtypes (e.g. Q1 through Q5) are denoted by color intensity.

Martini3 beads come in three sizes (regular, small, or tiny) and varying interaction affinities: bead types include hydrophobic, neutral, polar, charged, water, and halo-compounds (type C, N, P, Q, W and X, respectively; Figure 3D) (106). Interactions are tuned through bead subtypes, which indicate degree of polarity (1 being lowest), and chemical labels like hydrogen bond donor/acceptor (suffixes *a*/*d*). Additional flexibility is encoded by self-interaction labels “*h*” or “*r*”, which define higher or reduced self-interaction strength. We reasoned that scrambling should be determined primarily by interactions between protein beads and POPC, which has hydrophobic tails (types C1 and C4h), an intermediate/neutral glycerol backbone (types N4a, sN4a), and a charged head group (types Q1, Q5) (Figure 3E). Martini3 interactions are modeled as Lennard-Jones potentials; the interaction energies *εij* for all beads with POPC bead types vary significantly (Figure 3F). We hypothesized that a simple scramblase would demonstrate scrambling activity if it were constructed from beads with protein-like interaction profiles with POPC.

To identify which bead types are most suitable for a simple model scramblase, we conducted an initial survey of bead types. To do this, we converted all protein beads in the nhTMEM16-based scramblase CG model to a single, uniform bead type (iterating over all possible bead types) while retaining the shape of open-state nhTMEM16 with positional restraints (Figure 4A–E, Table S2). As a control, we simulated a position-restrained wildtype (WTpr) system with bead types determined by the default Martini3 mapping (Table S3), which matches the set of beads used for our flexible model of open state nhTMEM16. Compared to the flexible model with a scrambling rate of 12.3 ± 1.6 lipids⋅μs^−1^ (Figure 1E), WTpr models scramble 3.5 ± 0.7 lipids⋅μs^−1^, reflecting the contribution of small conformational rearrangements to scrambling.

**Figure 4.**
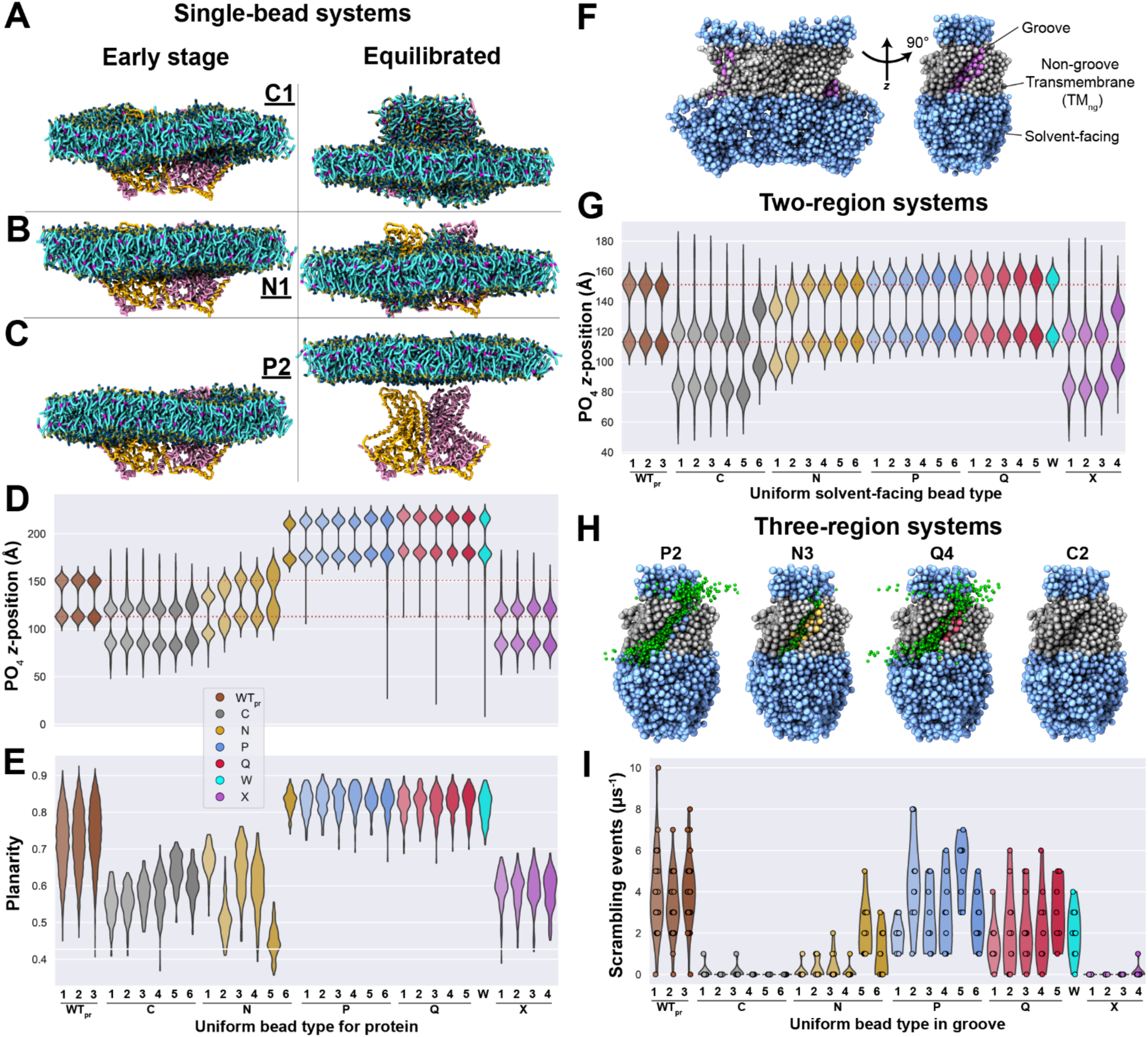
Selecting optimal Martini3 beads to model simple scramblases. (*A–C*) Initial (left) and final (right) lipid distributions for simple scramblases comprised of a single uniform bead type. Proteins are shown as in Figure 1, while POPC is cyan, red, blue, and tan. (*D*) Distributions of lipid head group *z*-positions for single-bead models. Red dotted lines indicate starting head group *z*-positions. (*E*) Lipid planarity distributions for single-bead models (see Methods). Values closest to one are most planar. (*F*) Illustration of bead type assignment: solvent-facing (sky blue), TMng (gray), and groove (pink) regions. (*G*) Head group distributions with bead C4 in TMng/groove regions but a varied bead in the solvent-facing region. (*H*) Three-component models with C4 TMng beads, P2 solvent-facing, and a changing bead type in the groove. Green spheres mark head group positions during scrambling events and labels denote groove beads. (*I*) Scrambling rates for three-region models. Models in (*A*–*C*) appear in Figure S4, and models in (*F*, *H*) appear in Figure S7. Data for the same WTpr system is shown in (*D*) and (*G*) and appears in Figures 5, S5-8, and S10.

For all uniform bead types tested, the membrane initially adopted a bilayer configuration (Figure 4A–C, left). When the protein was comprised of bead types with low polarity (e.g. C1), some lipids left the bilayer and coated all surfaces of the protein, including regions that should be solvent-exposed (Figure 4A, right; S4). Intermediate-polarity protein beads like N1 caused minimal distortion of the bilayer (Figure 4B, right), as reflected in the *z*-positions of the head groups (Figure 4D) and planarity of the membrane (Figure 4E) throughout the final eight microseconds of simulation. If the uniform protein bead was highly polar (e.g. P2), protein-water and protein-ion interactions displaced protein-lipid contacts and caused the bilayer to drift away from the protein (Figure 4C, right). We detected scrambling for N-, P-, and Q-type beads (Figure S5), with most of these events occurring before the bilayer moved away from the protein. Using a single uniform bead to represent the protein sometimes captured the head group *z*-distributions observed in WT_pr_ (Figure 4D) but did not recapitulate the planarity distributions in WTpr (Figure 4E). In most cases where scrambling was detected, passage of lipids did not occur through the canonical groove (Figure S5), so we concluded that a single-bead model cannot fully represent WTpr-like lipid scrambling.

### Constructing simple CG scramblases with distinct domains

To refine our simple scramblase model, we planned to partition it into non-groove TM (TMng), solvent-facing, and groove domains (Figure 4F), expecting this to yield more realistic scrambling than observed for uniform bead systems. Before moving to systems with unique beads in all three regions, we tested two-region models with unique beads in the TMall (including TMng and groove) and solvent-facing regions. For the TMall region, we selected bead C4 due to its low scrambling rate (expected for a TM region) in a mono-bead setup (Figure S5) and appearance in standard Martini3 definitions of Phe, Tyr, and Trp (106), residues commonly enriched in TM domains. Using domain boundaries from PPM 3.0 (112), we set TMall beads to type C4 and varied the solvent-facing bead types (Figure 4F, Table S4). Most N-, P-, and Q-type beads in the solvent-facing region resulted in head group distributions matching WTpr (Figure 4G) with little or no scrambling (Figure S6A–C). Among these possible beads, we chose type P2 beads to represent the solvent-facing regions due to its identity as the Martini3 backbone bead and the presence of P-type beads in topologies of Asn, Gln, Thr, and Ser (106). With this, we established a reasonable two-region model scramblase.

After selecting bead types for the TMall (hydrophobic C4) and solvent-facing (hydrophilic P2) regions, we divided the TMall region into independent TMng and groove regions, giving a total of three unique regions. Forty-eight beads were selected by inspection along the surface of the putative groove to define the groove region (Figure 4H, S7; Table S5). After varying the groove properties by assigning all groove beads uniformly to a single bead type, scrambling occurred primarily through the defined groove (Figure S7A), and selecting bead types N5–N6, P1–P6, Q1–Q5, or W resulted in a scrambling rate (Figure 4I) and time per event (Figure S7B) comparable to WTpr; this suggested that grooves spanning a range of interaction strengths with POPC (Figure 3F) scramble like WTpr. Since a P2 groove led to scrambling rates most consistent with WTpr (4.6 ± 1.6 lipids⋅μs^−1^; Figure 4I), we constructed our final calibrated scramblases with hydrophilic P2 beads in the groove and solvent-facing regions, and with hydrophobic C4 beads in the TMng region—we refer to this system as P23 (as it has 3 regions including a P2 pore). Although we defined three unique regions, the above process led us to assign the same P2 bead type to both the groove and solvent-facing regions; thus, P23 has only two unique bead types. Since the P23 model scrambles like WTpr but with significantly reduced complexity of surface beads, we used it as a baseline to examine other properties that are likely to affect scrambling.

To isolate the effect of modifying TMng beads without also changing groove-lining and solvent-facing beads, we converted all TMng residues in P23 back to bead types that match the nhTMEM16 sequence. This model, with a native-like TMng region but uniform bead types in the solvent-facing and groove domains (P2native), exhibited a scrambling rate of 3.4 ± 2.0 lipids⋅μs^−1^ (Figure S8; Table S3, S6), nearly unchanged from P23. This observation implies that the exact pattern of hydrophilicity in the TMng region resulting from the native sequence is not a key determinant of scrambling. Similarly, since P2native is like WTpr but with groove and solvent-facing beads set to a uniform type, the similar scrambling rate we observe for the two models suggests that the detailed hydrophilic pattern in the groove and solvent-facing regions is not a key determinant of scrambling. Therefore, scrambling can occur if the general hydrophilic nature of each region is similar enough to what is encoded by the wildtype sequence.

To test how the surface geometry of residues in the groove affects scrambling, we further modified the P23 model by assigning uniform alanine shapes to all residues, including the 48 residues that form the groove, without changing their bead types (P2polyala) (Figure 5A, Table S7). This changed the surface geometry, but not the hydrophilicity in the groove or its overall architecture. The modified groove in P2polyala scrambled at 0.46 ± 0.35 lipids⋅μs^−1^ (Figure 5B), roughly sevenfold reduced compared to P23, suggesting that surface geometry is important, but not essential for scrambling.

**Figure 5.**
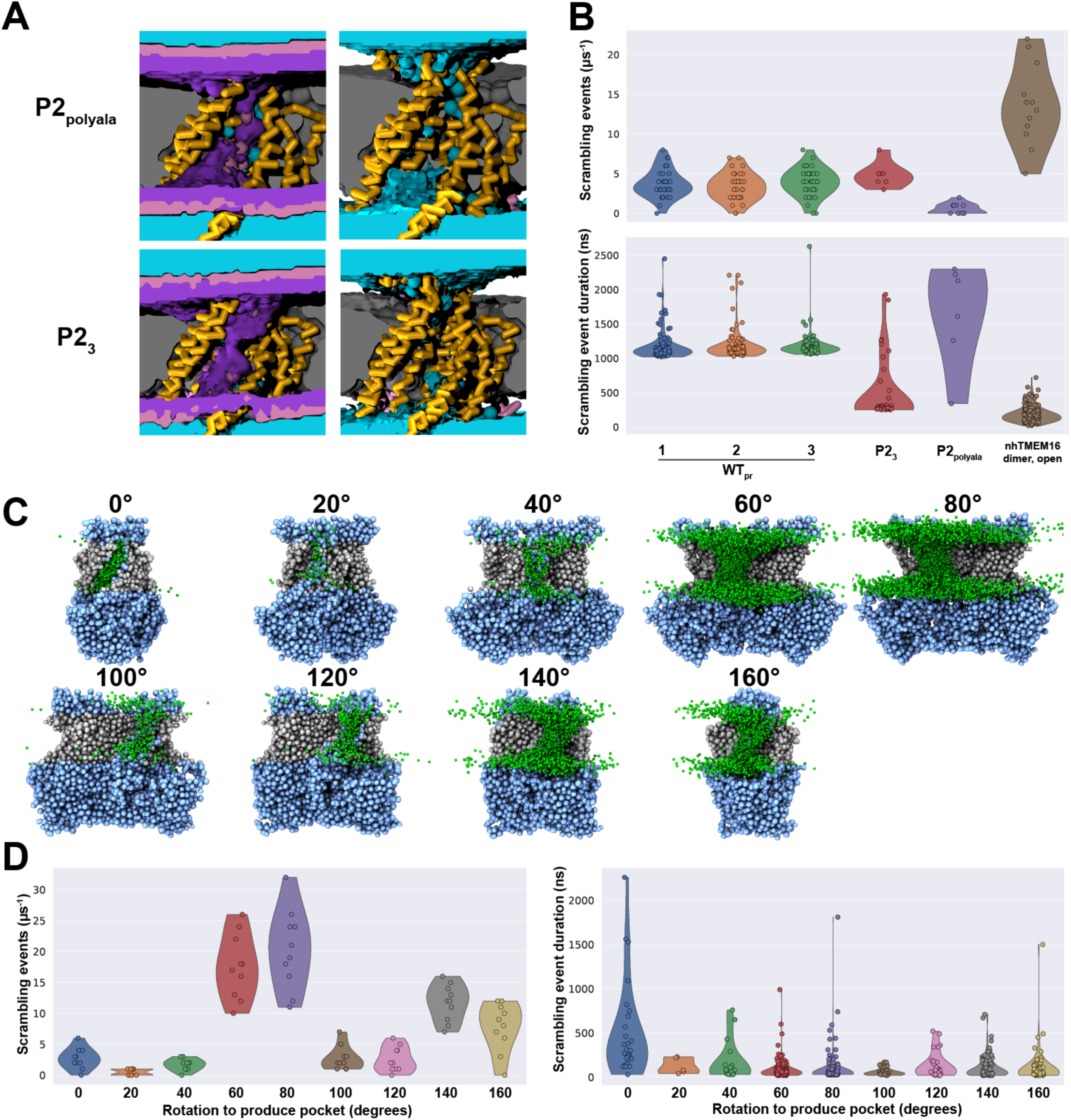
Groove architecture controls scrambling activity in simple models. (*A*) Densities for water (cyan), lipid head groups (purple), and overlap between the two (pink). nhTMEM16 is shown in pink and yellow cartoon. (*B*) Scrambling rate (top) and duration per event (bottom) for the P2_polyala_ model compared to position-restrained WT_pr_, P2_3_, and the flexible model with elastic network bonds. (*C*) Scrambling by three-region scramblases after rotating the groove about the *z*-axis by the indicated angle (see Methods and Movie S10). In *C*, C4 TMng beads are gray and P2 solvent-facing/groove beads are sky blue while green spheres mark head group positions during scrambling. (*D*) Scrambling rate (*left*) and time per event (*right*) for three-region scramblases after groove rotation. Data in (*B*) also appear in Figure 1 (flexible nhTMEM16 model) and Figure 4I (P23 and WTpr). Data for WTpr appears in Figures 4, 5, S5-8, and S10.

### Scrambling through grooves with varied architecture

Having established that the structural architecture common to TCS proteins facilitates scrambling when using groove-lining beads with differing interaction propensities and surface geometries, we speculated that changing the architecture of the groove might have a significant influence on scrambling rates. We generated protein models with groove-like patches, i.e., with a patch having similar surface area and polarity to the original groove. To do so we rotated groove beads about the center of the protein and projected them onto its surface (Table S8; see Methods). Our procedure generated hypothetical groove-like patches (made of P2 beads) at different locations around the membrane-exposed surface of nhTMEM16, but not at the putative scrambling-active sites. Generated groove-like patches inherited the architecture of TM-exposed portions of nhTMEM16, so these models can address whether the distinct architecture of the native groove is a prerequisite to scrambling activity.

Model scramblases generated based on groove rotations of 20, 40, 100, and 120° resulted in models with groove-like patches that scrambled at levels near the native groove placement (0.5 to 2.6 lipids⋅μs^−1^), whereas groove rotations of 60, 80, 140, and 160° led to scrambling rates from 11.7 to 20.3 lipids⋅μs^−1^ (Figure 5C, D and Movie S10). Groove rotations leading to higher scrambling rates tended to position the groove-like patch on an outcropping. In contrast to the invagination of the conventional groove, outcroppings can accommodate multiple lipid head groups at a given distance through the bilayer, which grants translocating lipids more configurational freedom than available in the native groove; this may explain the observed increase in scrambling rates for some groove-like patch architectures. All alternative architectures tested led to scrambling events of shorter average duration (60–180 ns) than what was observed for the canonical groove (511.1 ± 210.6 ns; Figure 5D), suggesting that the canonical groove has an architecture tuned to restrict scrambling rates. Unlike our hypothetical groove-like patch architectures, the invagination of the canonical groove offers control over the degree of openness and therefore the exposure of hydrophilic groove residues. Control over scrambling activity likely allows scramblases to be activated specifically during signaling events.

### Scrambling activity is elevated in scramblases with thinner TM**ng** regions

Scramblases may thin the membrane to promote scrambling (13, 59, 69, 76, 88, 91–93). Some scramblases induce more membrane thinning than others in MD simulations (91), so we reasoned that the degree of thinning should be a function of how hydrophobicity is distributed in the TM region. We returned to our original P23 model scramblase and systematically varied the thickness of the TMng region (Figure 6A, B, Table S9). For our thinnest TMng region (thickness: 10 Å), we recorded robust scrambling at 17.8 ± 3.4 lipids⋅μs^−1^, elevated from the 2.3 ± 0.9 lipids⋅μs^−1^ for native TMng region thickness (34 Å). Only three other systems exhibited an elevated scrambling rate compared to the native width: TMng region thicknesses of 14, 18, or 22 Å led to scrambling at 8.2 ± 2.4, 3.1 ± 1.1, and 3.6 ± 1.1 lipids⋅μs^−1^, respectively.

**Figure 6.**
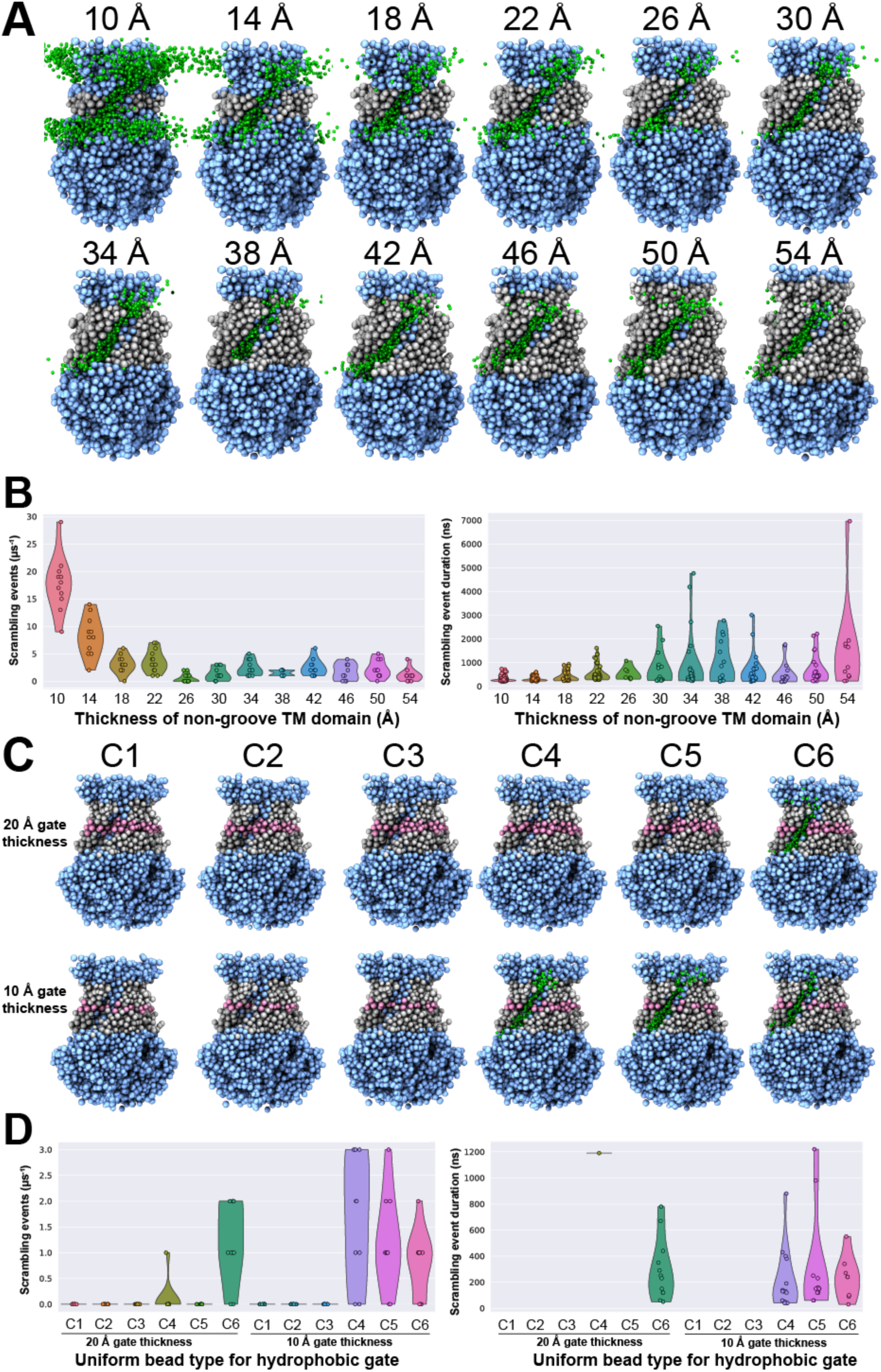
Thickness of hydrophobic transmembrane regions and hydrophobic gates can control scrambling. (*A*) Systems are labeled by the approximate thickness of the TMng region (gray C4 beads). Groove and solvent-facing beads are type P2 (sky blue). (*B*) Scrambling rate (*left*) and duration per event (*right*). Hydrophobic gates decrease scrambling rate. (*C*) Scramblases with hydrophobic gates (pink) of thickness 20 Å (*thick*) or 10 Å (*thin*) and varied bead type (C1–C6). (*D*) Rate (*left*) and duration (*right*) of scrambling for each hydrophobic gate.

Although the energy barrier opposing scrambling is decreased in our models with thinner TMng regions, scrambling occurred only through the groove formed by α3–α7 (Figure 6A). TMng regions of widths from 26 to 54 Å exhibited similar scrambling rates (0.5 to 2.5 lipids⋅μs^−1^), perhaps because the energetic barrier to scrambling in these cases is determined by the intrinsic width of the bilayer. The time per scrambling event increased slightly as a function of TMng region thickness: events for TMng regions thicker than 30 Å lasted 256–1200 ns⋅event^−1^ on average while we observed events lasting 126.1 ± 14.2 ns⋅event^−1^ in the thinnest TMng region (Figure 6B, right). These results suggest that the thickness of the TMng region directly controls both scrambling rate and time per event.

### Gates of varying hydrophobicity along the groove block scrambling

Three hydrophobic residues in the groove of mmTMEM16F form an inner activation gate that inhibits scrambling (87), and the equivalent of a single hydrophobic-to-polar mutation in α4 (e.g. F518K in mmTMEM16F) results in a constitutively active scramblase for several TCS proteins (37). We hypothesized that disrupting the hydrophobic gate in TCS proteins could stabilize the scrambling-competent open state and it could simultaneously decrease the energetic barrier to scrambling once the open state is reached. Since our simple scramblases based on nhTMEM16 lack a hydrophobic gate, we added hydrophobic gates with varied width and hydrophobicity and recorded scrambling activity (Figure 6C, D; Table S10).

Gates of ∼20 Å thickness were sufficient to block scrambling for hydrophobic bead types C1–C5, but we observed scrambling at 1.2 ± 0.8 lipids⋅μs^−1^ for the most hydrophilic C-type bead, C6 (Figure 6D, left); this is decreased compared to systems without a hydrophobic gate (P23, 4.6 ± 1.6 lipids⋅μs^−1^; Figure 4I). To put the hydrophobicity of C6 beads into context, C6 beads are found in Martini3 definitions of Cys and Met side chains. For thinner gates of ∼10 Å thickness, scrambling did not occur if gates were comprised of lower polarity C-type beads (C1–C3), but higher polarity C4–C6 gates permitted scrambling at 0.6 to 1.2 lipids⋅μs^−1^. Therefore, hydrophobic clusters interrupting the groove can block scrambling directly, not only by favoring the closed state configuration. Surprisingly, we observed that the time per scrambling event for all scrambling-competent systems with a hydrophobic gate, 210 to 304 ns⋅event^−1^ (Figure 6D, right), was slightly reduced compared to the 470 ± 120 ns⋅event^−1^ for P23. We hypothesize that the time per scrambling event decreases in cases where interactions between the pocket and lipid are less favorable. These simulations illustrate that hydrophobic patches interrupting the groove both decrease scrambling rate and decrease the time per scrambling event, highlighting the importance of net groove hydrophilicity.

### Scrambling through simple grooves with well-defined basic shapes

Given that our nhTMEM16-based models indicate that groove architecture might be a key determinant of scrambling over groove hydrophilicity and surface geometry, we wondered whether even simpler models could promote scrambling. Inspired by simple models used to test constant electric fields in MD simulations (127), we constructed scramblases with a cubic architecture of similar size to our protein-based models (side length 80 Å). We partitioned the cubic scramblases into TMng, solvent-facing, and groove regions with the same bead types and region sizes as P23 models and embedded them in POPC bilayers (Figure 7A, top). All “protein” beads in the cube had the same size. Using standard definitions of shapes (spheres, rectangles, cylinders, and ellipsoids) we created customized grooves by adding or removing beads on each membrane-adjacent cubic face (Figure 7A, S9) and simulated each model for at least 10 μs (Table S11). A minority of grooves permitted enough water to flow through them that the semiisotropic barostat led to instabilities; in these cases, we switched to an isotropic barostat and reduced the timestep (Table S12). Usage of the isotropic instead of the semiisotropic barostat did not affect scrambling for P23 (Figure S10).

**Figure 7.**
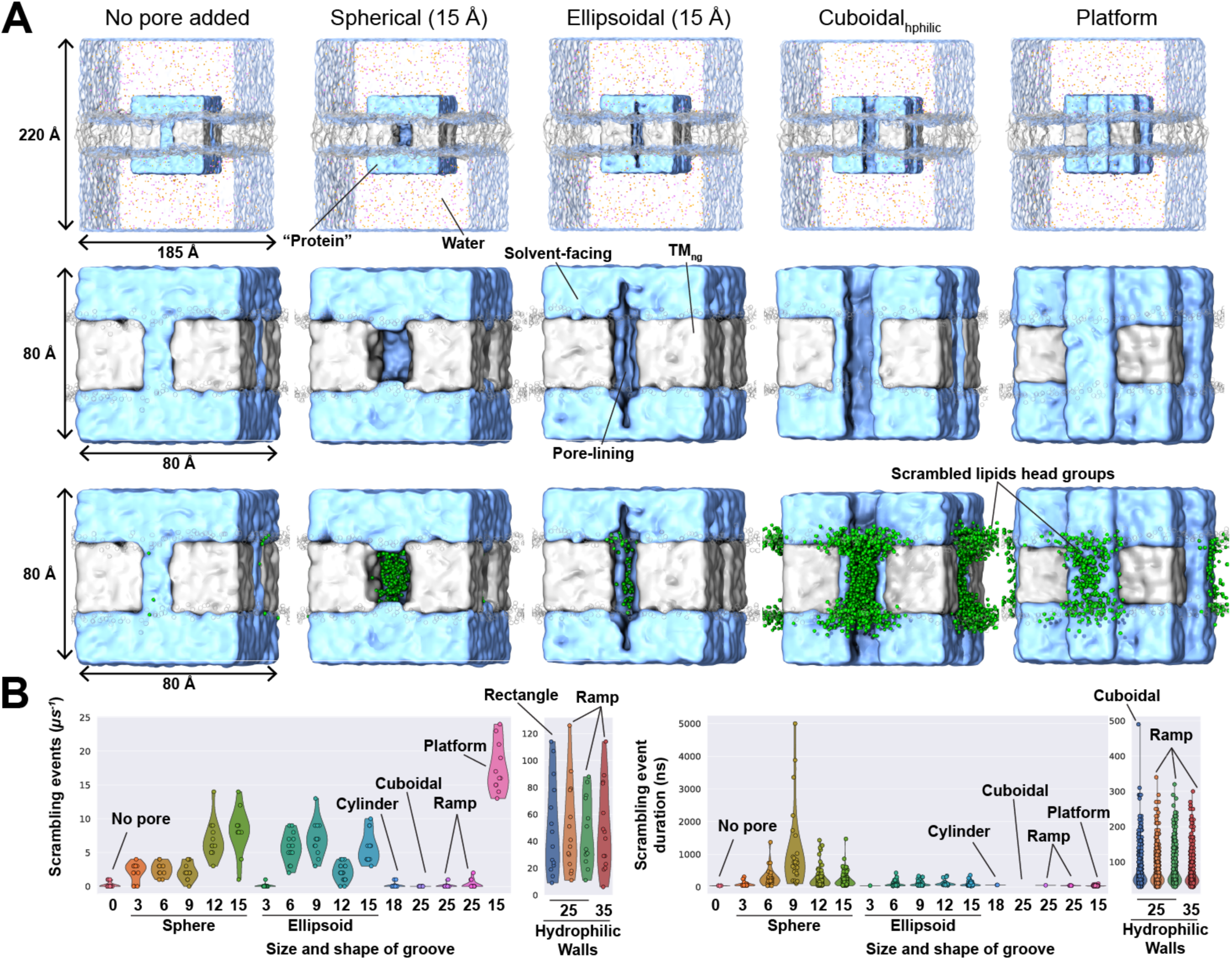
Simple scramblase models. (*A*) Overall system setup (*top*), design of the groove (*middle*), and scrambling (*bottom*) by cubic scramblases with grooves of the indicated shape on each TM face. Hydrophilic beads (blue, type P2) and TMng beads (gray, type C4) are shown. The lengths in parentheses are the shape factors, which control the size of each groove (larger shape factors indicate larger grooves). The subscript *hphilic* reflects that hydrophilic P2 beads were used to form all walls of a groove. Green spheres mark head group positions during scrambling. In (*A*, *top*), water is transparent blue while positively and negatively charged ions are pink and orange, respectively. (*B*) Scrambling rates (*left*) and time per event (*right*) for scrambling by cubic models. Numeric labels on the *x*-axes are shape factors. Models in (*A*) appear in Figure S9.

For a purely cubic scramblase with polar beads forming a flat pathway spanning the TM region (no pore, groove or outcropping), only 0.2 ± 0.2 lipids⋅μs^−1^ crossed the membrane (Figure 7A, B). This implies that some architectural features of a groove facilitate scrambling, e.g., by providing space for polar head groups to minimize interactions with hydrophobic tails during translocation. Indeed, spherical grooves enabled scrambling: spheres of radius 3, 6, and 9 Å scrambled 2.2 to 2.5 lipids⋅μs^−1^ while those of radius 12 and 15 Å scrambled 6.8 ± 1.7 and 8.2 ± 2.1 lipids⋅μs^−1^, respectively (Figure 7B, S9 and Movie S11). With increasing radius, spheres form grooves of greater depth that span a larger portion of the TM region; both factors may facilitate scrambling.

To evaluate the importance of groove depth, we constructed ellipsoidal grooves that span the full *z*-extent of the cubic scramblase but vary in maximal groove depth (Figure S9). The shallowest ellipsoid (shape factor: 3 Å) scrambled 0.1 ± 0.2 lipids⋅μs^−1^, whereas deeper ellipsoids (shape factor: 6 to 15 Å) scrambled from 2.1 to 5.8 lipids⋅μs^−1^ (Figure 7B). Therefore, membrane-spanning grooves appear to scramble if they reach a threshold depth (∼6 Å), with little change to scrambling rate for depths greater than the threshold (Figure 7B). This threshold depth effect likely arises because some shielding of hydrophilic head groups from nonpolar lipid tails is conducive to scrambling, but lipids do not penetrate enough to reach the innermost wall of the groove.

Given that outcroppings were associated with increased scrambling in our nhTMEM16-based models with hypothetical, rotation-based “grooves” (Figure 5C, D), we constructed a cubic scramblase with cuboidal outcroppings extending three beads (15 Å) above the cubic faces of the scramblase (*platform* system; Figure 7A, Movie S12). These platforms scrambled lipids at a net rate of 17.5 ± 2.5 lipids⋅μs^−1^ (Figure 7B, left), equivalent to 4.4 lipids⋅μs^−1^⋅groove^−1^, which is slightly less than the rates observed for rotation-based outcroppings (5.9 to 10.2 lipids⋅μs^−1^⋅groove^−1^). The rough surfaces of protein-like outcroppings may lead to higher scrambling rates than flat cuboidal outcroppings because changes in depth along the scrambling pathway on a protein-like surface can occur incrementally instead of concomitantly.

Not all membrane-spanning shapes led to scrambling in our simulations—cylindrical, cuboidal, and ramp geometries had net scrambling rates between 0.1 to 0.3 lipids⋅μs^−1^. Unlike the ellipsoidal grooves, these four non-scrambling geometries featured hydrophobic groove walls, so we built four new cuboidal-like systems with hydrophilic (hphilic) P2 walls: cuboidalhphilic, downward ramphphilic, stepped ramphphilic, and curved ramphphilic (Figure S9), which scrambled 46.7 to 50.3 lipids⋅μs^−1^ (Figure 7B). These efficiently scrambling, hydrophilic grooves differed in their depth profiles across the bilayer, but this minimally affected scrambling rates. Interestingly, the sharp corners of cuboidal grooves did not preclude scrambling compared to the smoother edges of ellipsoidal and spherical grooves. The higher scrambling rate for cuboidal compared to ellipsoidal systems may be attributable to the larger groove width for cuboidal geometries. Taken together, a comparison of grooves with or without hydrophilic pore-lining walls reveals that a fully hydrophilic groove allows substantially more scrambling than only partially hydrophilic grooves.

Overall, scrambling events happened quickly for simple grooves (∼60–200 ns; Figure 7B right) compared to WTpr (∼800 ns; Figure S7B, left), except for systems with spherical grooves where events lasted from 200 to 1,100 ns per event on average (Figure 7B, right). This implies that groove architecture alone can modulate the duration per scrambling event. Additionally, groove architecture strongly affected scrambling rates, and several shapes prevented scrambling entirely. However, numerous unique groove architectures led to scrambling rates on the same order of magnitude, demonstrating that a variety of unique groove shapes can scramble effectively. In summary, our results from simulations of simple systems predict that scrambling rates are maximized when the groove is of sufficient depth, the groove is of sufficient width, and the groove walls are hydrophilic. These minimal restrictions are encouraging for *de novo* scramblase design.

## DISCUSSION

Scramblases are key players in critical biological processes such as blood coagulation (9), lipid synthesis (5, 6, 8), apoptosis (17, 36), and autophagy (10, 33). Emerging evidence suggests that the capacity to scramble may be conserved across TCS family members (37, 59), which may implicate scrambling in inner-ear sensory cell function (TMCs) (49, 94, 95), neuronal development and humidity sensing (TMEM63s) (128–130), or drought detection in plants (OSCAs) (52, 56). The role of lipid scrambling in these processes is currently unclear. Additionally, there are substantial sequence differences across TCS proteins, so the shared factors leading to scrambling activity are difficult to systematically evaluate. Consistent with the credit card mechanism for scrambling (47, 76, 84), the results of our CG simulations suggest that scrambling activity is largely determined by the architecture of the groove and fine-tuned by the hydrophilicity and surface geometry of groove-lining residues.

Our CG simulations of simple systems show that scrambling rates are maximized in wide, deep grooves with hydrophilic residues lining all walls. Raised hydrophilic surfaces in nhTEM16-based models with rotated “grooves” and in cubic models also scrambled efficiently. Notably, these are not the architectures adopted by TCS proteins; this indicates that the characteristic invaginated groove adopted by TCS family members is not optimized to scramble efficiently, but rather to offer conformational control of scrambling activity. Relatedly, we observed a decrease in scrambling rate and time per event after introducing hydrophobic gates in simple nhTMEM16-based models. Since some TCS proteins have hydrophobic gates in the groove, this appears to be a control mechanism to prevent or slow unwanted scrambling in the open state. We speculate that the hydrophobic gate may also energetically favor the closed state, preventing scrambling by keeping the groove closed.

Interestingly, we observed membrane curvature changes per leaflet, which extended away from the simulated proteins, but we did not observe substantial membrane thinning in our scrambling-competent models except near the groove (Figure S3). This local thinning by non-scrambling lipids may be a consequence or a cause of lipid scrambling, and might depend on coarse-graining procedure, membrane composition, and the simulation timescale. Regardless, local membrane thinning was more apparent for open-like than for closed grooves (Figure S3) and might be correlated with pore water occupancy. We observed little closed-groove scrambling in all protein-based systems but one–CG simulations of hsTMC1 in its presumed closed conformations (with and without its essential hsCIB2 cytoplasmic partner) did reveal scrambling, although at rates that were significantly lower than those monitored for the presumed open state. It is possible that our TMC1 models, which are based on experimentally validated AF2 and AF3 predictions (45, 46, 48, 94), are not capturing true closed states. On the other hand, TMC proteins might have a basal scrambling activity enhanced by membrane tension/deformation. In a physiological context, this basal scrambling activity may be suppressed by other binding partners or the properties of the membrane.

All our CG simulations of nhTMEM16, hsTMEM63B, and atOSCA1.2 in closed and open states use experimentally derived structural models. For hsTMEM63B and atOSCA1.2, only the open-like states scrambled lipids. Simulations for the closed state of nhTMEM16 showed greatly reduced scrambling, consistent with evidence that closed-state nhTMEM16 exhibits some scrambling activity (69). Given that many members of the OSCA/TMEM63 family are mechanosensitive ion channels activated by membrane stretch (51, 53–55, 61, 64), our results suggests that their scrambling activity is controlled by mechanical stimuli.

Open-state cryo-EM structures of mechanosensitive OSCA/TMEM63 proteins have been obtained in highly curved liposomes (53), after changing membrane properties by adding methyl-beta-cyclodextrin (53), or by mutating residues in the hydrophobic gate (59). Scrambling in TMEM63B can be triggered by removal of cholesterol or lipid cleavage by extracellular phospholipases (59). Overall, it seems that changes to the structural properties of the membrane activate these scramblases. However, our simulations are in pure POPC with no applied surface tension, which may change the mechanics of scrambling by underestimating membrane thickness (131) and overestimating fluidity (132). Protein conformational changes are not described in our Martini3 CG models, but all-atom MD simulations could conceivably model the closed-to-open transition and lipid scrambling if the correct stimulus is applied over sufficiently long timescales.

Most of the scrambling events in our protein-based models occurred with lipid head-groups going through or on the surface of the hydrated groove. These results suggest that lipids head groups do not need to fully insert into the groove to be flipped (“dry” credit card mechanism)—in many instances, a hydrated pore favors the passage of the lipid head groups on or near the surface of the open groove (“wet” credit card mechanism). This may explain lack of lipid specificity and suggests why precise protein-lipid interactions may not be needed for lipid scrambling (13, 92). In addition, we detected low-probability scrambling across alternative, noncanonical pathways in some TCS proteins (Movie S2, S5, S7). Scrambling through these noncanonical pathways may depend on the hydrophobicity of the surfaces involved. Overall, our results support dry and wet credit card mechanisms (84) but cannot rule out a subtle role for noncanonical and closed-groove scrambling pathways.

We recorded scrambling rates up to 6.2⋅10^6^ lipids⋅s^−1^⋅groove^−1^ in Martini3 models of TCS family members and up to 6.9⋅10^6^ lipids⋅s^−1^⋅groove^−1^ in simple cubic models. Experimentally, scrambling rates have not been determined for many TCS proteins, but scrambling rates of 7.1⋅10^4^ lipids⋅s^−1^ for mmTMEM16F (103) and >2⋅10^4^ lipids⋅s^−1^ for afTMEM16 have been reported (92). One of the fastest known scramblases, opsin, scrambles >10^5^ lipids⋅s^−1^ (92). Therefore, we overestimate scrambling by about two orders of magnitude for TCS members and record rates one order of magnitude faster than opsin; this may be a limitation of Martini3 coarse-graining (91). Although Martini3 depicts molecular shape more accurately than Martini2 (106), Martini3 models scramble around 100-fold faster than Martini2 models of MTCH2 (31). Additionally, we use an elastic network model to maintain tertiary structure of simulated TCS family members, which allows fluctuations relative to the experimental structure; for nhTMEM16, the flexible model scrambles four times faster than the position-restrained model. Despite sampling limitations, all-atom simulations appear to match experimental scrambling rates more closely than CG Martini3 models (85–89, 93, 126), but simulations of mmTMEM16F lasting long enough to obtain good statistics (100 scrambling events) would have to reach ∼5 milliseconds of simulation time (assuming 2⋅10^4^ lipids⋅s^−1^) if rates are modeled correctly, and this is not easily accessible on modern computing hardware.

Besides the nature of the CG force field, our models of nhTMEM16 and hsTMC1 with hsCIB2 do not explicitly include bound Ca^2+^, although physiologically these should be present in the simulated open-like conformations of nhTMEM16. We observe similar rates to TMEM16 systems with DOPC and Ca^2+^ (91), suggesting that the lack of bound ions has a minor effect in our simulations, likely because we intentionally prevent closing of the open-like conformation with constraints. These results suggest that Ca^2+^ is only needed to stabilize the open-like groove in TMEM16 scramblases.

Our CG simulations do not include membrane voltage, so it is difficult to evaluate the pathway that ions would take in open conductive states of TCS proteins. Incorporating an applied biasing voltage using a non-polarizable CG force field is problematic, but we can infer the ion conduction pathway by inspecting equilibrium ion densities (Figure S11). These indicate that either positively or negatively charged ions occasionally visit the groove in open-like nhTMEM16 (Figure S11A) and hsTMEM63B (Figure S11B), that ions do not visit the groove of atOSCA1.2 as often (Figure S11C), and that hsTMC1 has a moderate preference for positively over negatively charged ions (Figure S11D, E). These apparent differences in ion preference likely depend on the membrane composition, which is pure POPC in our models. Despite the inherent limitations of CG models, our work suggests that TCS proteins may conduct ions through the open groove required for robust lipid scrambling. However, whether lipid headgroups and ions share the exact same translocation pathway or travel separately with lipids forming a pore wall remains to be elucidated.

The biophysical principles of lipids scrambling by TCS proteins and simple models uncovered here can guide design of *de novo* lipid scramblases. We suggest that groove architecture is a critical determinant of controlled lipid scrambling, which can be used to design proteins with substantially increased scrambling rates that may help uncover the elusive role of scrambling in physiological processes like auditory mechanotransduction (49, 94, 95).

## AUTHOR CONTRIBUTIONS

M.S., T.H.R., and H.E.S. designed the project. H.E.S. performed all simulations and data analyses. M.S. supervised all work and provided support through the entire project. M.S. verified all the methods mentioned in this manuscript. W.Z. and A.B. guided data interpretation. H.E.S., T.H.R, and M.S. wrote all sections of the manuscript with contributions from W.Z and A.B. All authors discussed the results and contributed to the final manuscript.

## ACKNOWLEDGMENT

Simulations were performed at the Ohio Supercomputing Center supercomputer Pitzer (grants PAS1037 and PAA0217), Texas Advanced Computing Center supercomputer Stampede3 and the San Diego Supercomputing Center supercomputer Expanse (ACCESS MCB140226). We acknowledge insightful discussions with Jeffrey R. Holt and members of the Sotomayor lab. We thank Wei-Hsiang Weng for sharing AF3 models of open-state hsTMC1 and Yaw Agyemang for providing AF2 models of hsTMC1 with and without hsCIB2.

## SUPPORTING CITATIONS

No references appear in the Supporting material.

## SUPPORTING INFORMATION

**Movie S1.** A scrambling event in open-like nhTMEM16 through the canonical pathway. Protein is shown in pink and orange surface, non-scrambling lipid head groups are transparent gray, and the scrambling lipid shown has its head group (name PO4) in magenta. All positions are shown every 10 ns with time-averaged positions over 30 ns.

**Movie S2.** A scrambling event in open-like nhTMEM16 through a noncanonical pathway. Shown as in Movie S1.

**Movie S3.** A scrambling event in open-like hsTMEM63B through the canonical pathway. Shown as in Movie S1.

**Movie S4.** A scrambling event in open-like atOSCA1.2 through the canonical pathway. Shown as in Movie S1.

**Movie S5.** A scrambling event in open-like atOSCA1.2 through a noncanonical pathway. Shown as in Movie S1.

**Movie S6.** A scrambling event in closed dimeric hsTMC1 with two bound hsCIB2s through the canonical pathway. Shown as in Movie S1.

**Movie S7.** A scrambling event in closed dimeric hsTMC1 with two bound hsCIB2s through a noncanonical pathway. Shown as in Movie S1.

**Movie S8.** A scrambling event in open-like dimeric hsTMC1 through the canonical pathway. Shown as in Movie S1.

**Movie S9.** A scrambling event in open-like dimeric hsTMC1 through a noncanonical pathway. Shown as in Movie S1.

**Movie S10.** A scrambling event in P23 after the groove has been rotated by 60°. Protein is shown in coarse surface and colored according to bead type (gray is hydrophobic, blue is hydrophilic). Non-scrambling lipid head groups are transparent gray, and the scrambling lipid shown has its head group (name PO4) in magenta. All positions are shown every 10 ns with time-averaged positions over 30 ns.

**Movie S11.** A scrambling event in the cubic model with a spherical groove shape (radius 15 Å). Shown as in Movie S10.

**Movie S12.** A scrambling event in the cubic model with a “platform” groove shape. Shown as in Movie S10.

**Figure S1.**
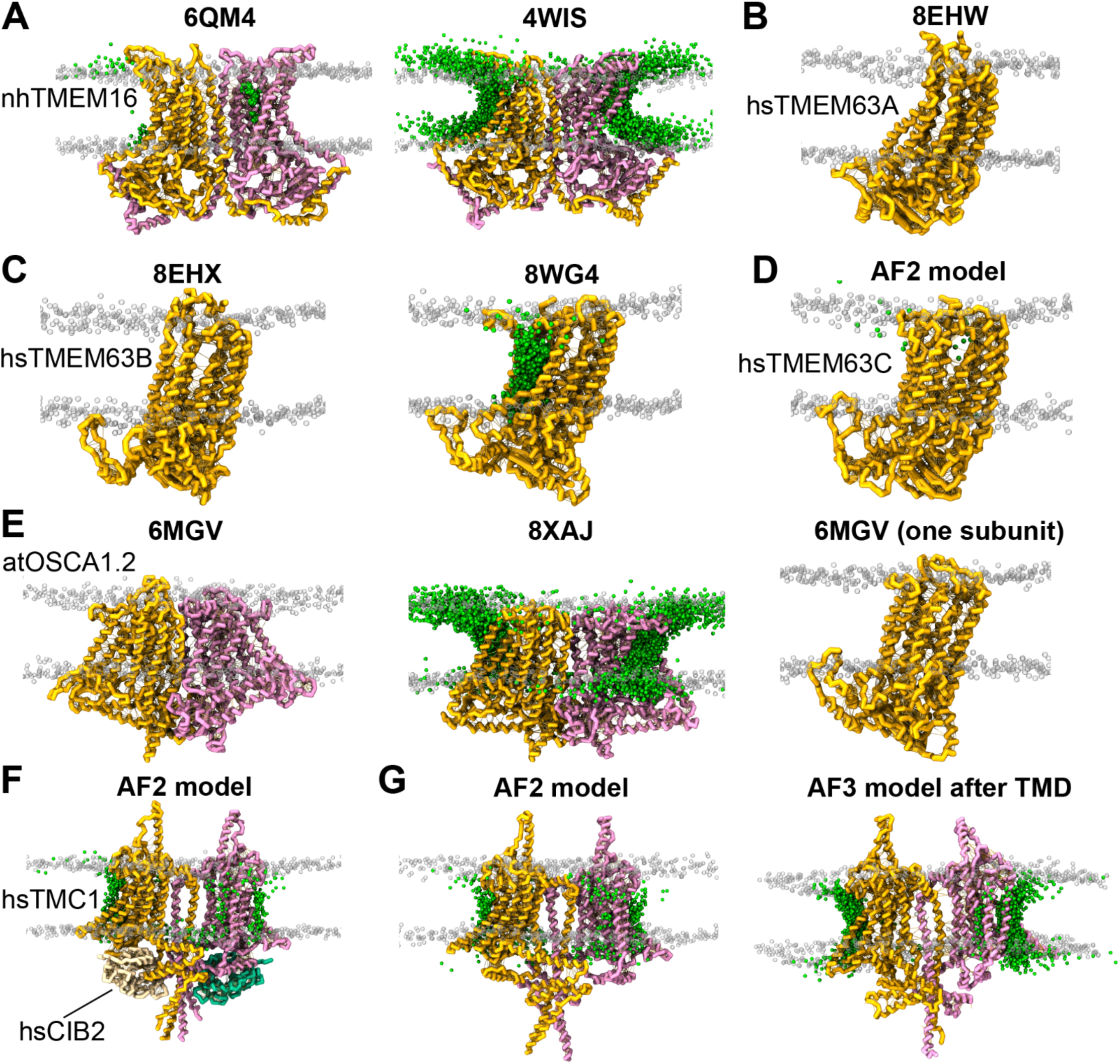
Scrambling for TCS family members using CG models. (*A*–*G*) Initial structures overlaid with head group paths taken throughout any scrambling events detected. Transparent gray spheres mark non-scrambling phosphate head groups and green spheres illustrate the path taken by lipid head groups during scrambling. Models from (*A*), (*C*), (*E*), and (*F*) appear in Figure 1.

**Figure S2.**
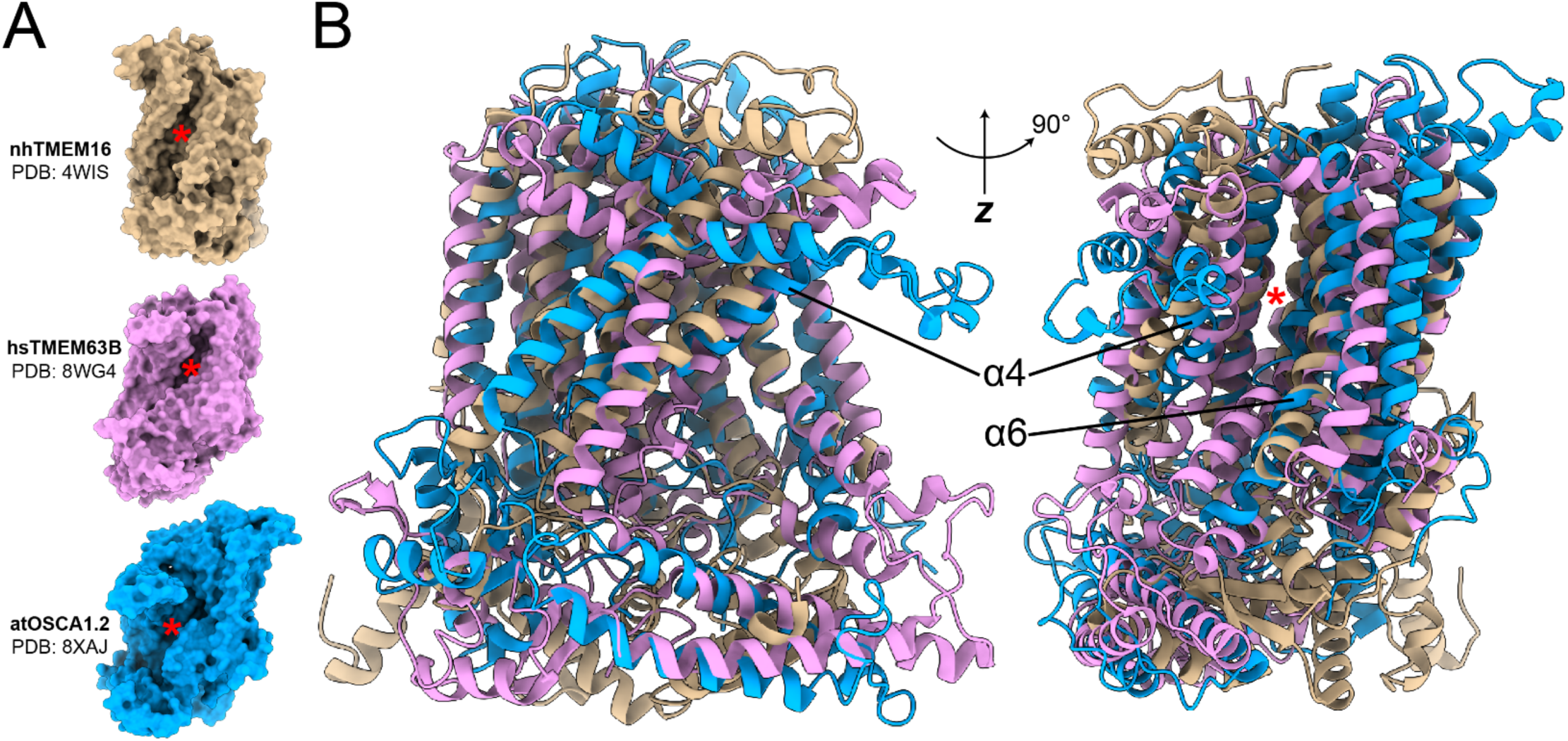
Comparison of groove architecture in open-state TCS protein models. (*A)* Surfaces are shown for crystal or cryo-EM structures of nhTMEM16 (tan; PDB: 4WIS), hsTMEM63B (pink; PDB: 8WG4), and atOSCA1.2 (blue; PDB: 8XAJ). Red asterisks mark the approximate locations of the grooves. (*B*) MatchMaker-aligned models rotated about the *z*-axis by 90° (*left*) and in the same orientation as (*A*) (*right*) to illustrate differences in separation between α4 and α6. Only one subunit is shown for dimeric nhTMEM16 and atOSCA1.2.

**Figure S3.**
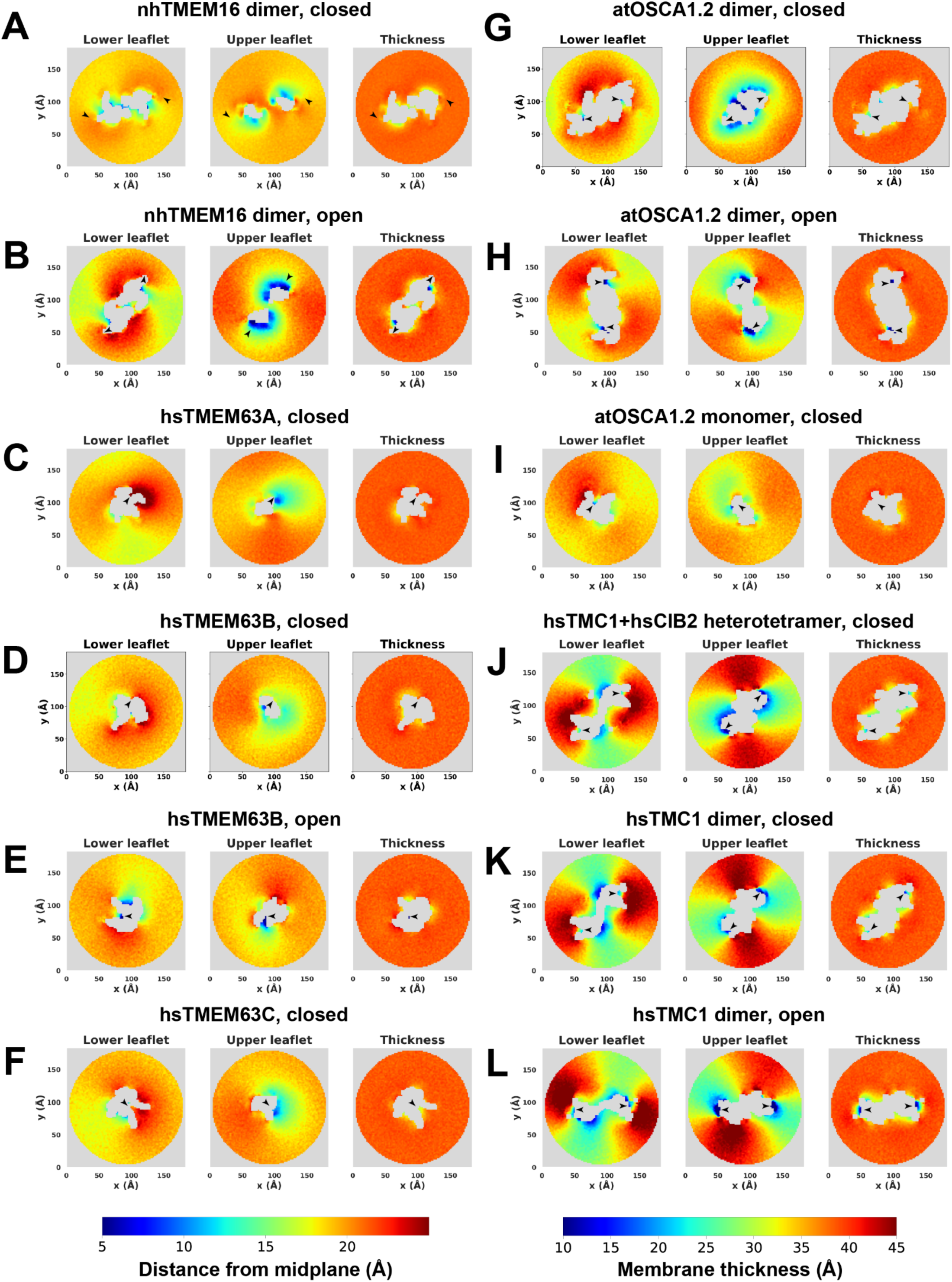
Membrane curvature and thickness in TCS family proteins. (*A*–*L)* 2D histograms of lipid head group (name PO4) *z*-positions in the lower *(left)* and upper *(middle)* leaflets averaged throughout each simulation. (*Right*) averaged membrane thickness over time, obtained as the difference between lower and upper leaflets. Color scales are shared for all plots; distance from midplane (*left)* applies to leaflet *z*-positions (*A*–*L*, *left* and *middle*) while membrane thickness (*right*) applies to membrane thickness (*A*–*L*, *right*). Arrowheads mark the approximate location of each groove.

**Figure S4.**
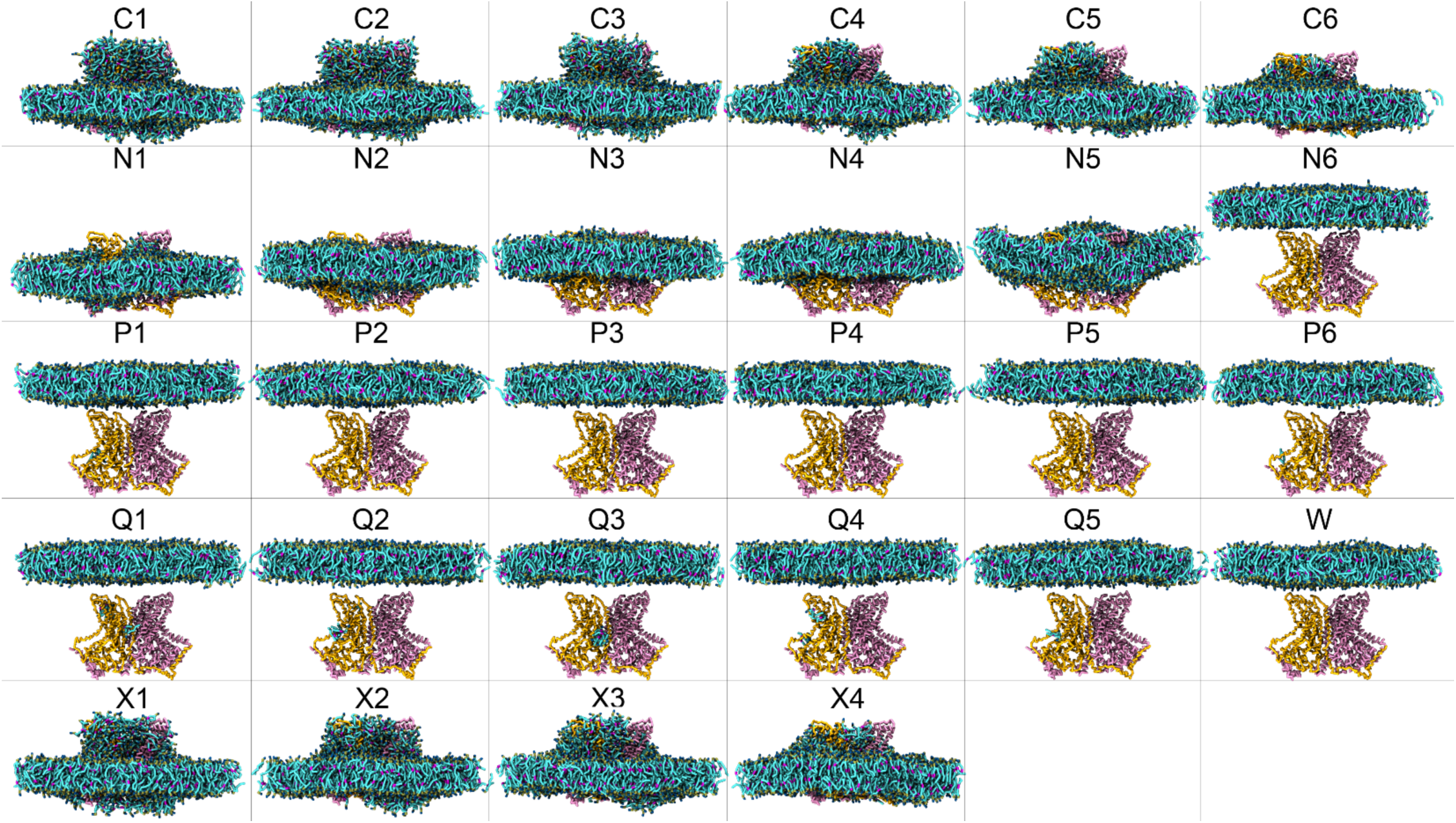
Final state reached in the simulations with protein represented as a uniform bead type. Protein is shown in orange and pink ribbons while POPC is cyan, red, blue, and tan. For hydrophobic beads, lipids also coat the protein (C1–C6; X1–X4). Hydrophilic protein beads interact more favorably with water and solvent (N6; P1–P6; Q1–Q5; W). Protein beads with more balanced interactions allow the bilayer to remain intact around the protein TM domains (N1–N5). Models for C1, N1, and P2 appear in Figure 4.

**Figure S5.**
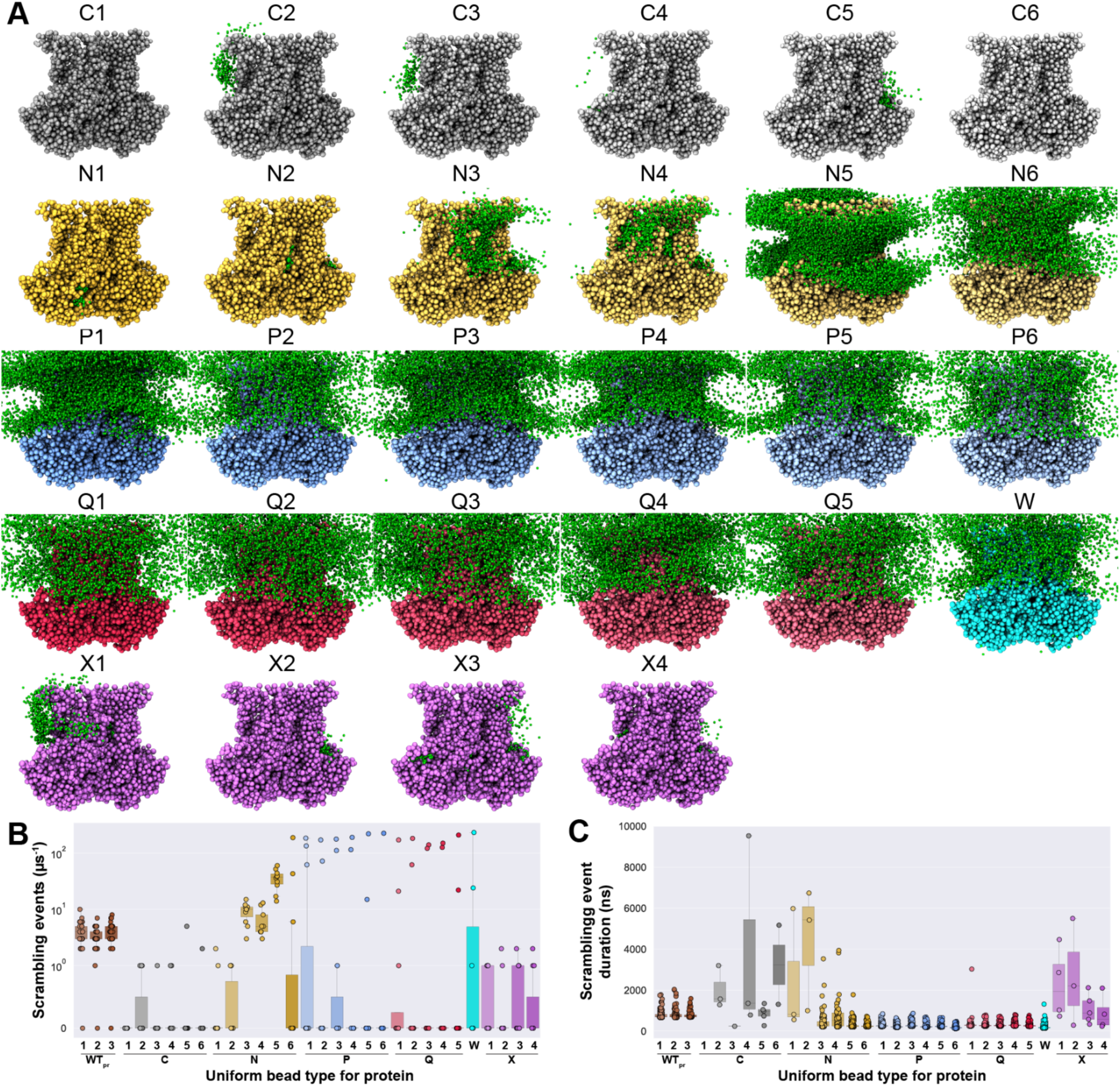
Scrambling by single-bead systems with shape based on dimeric nhTMEM16. (*A*) Systems after an initial minimization and equilibration. Green spheres are head group positions during scrambling events. (*B*, *C*) Scrambling rate and duration per event. Data for WTpr appears in Figures 4, 5, S5-8, and S10.

**Figure S6.**
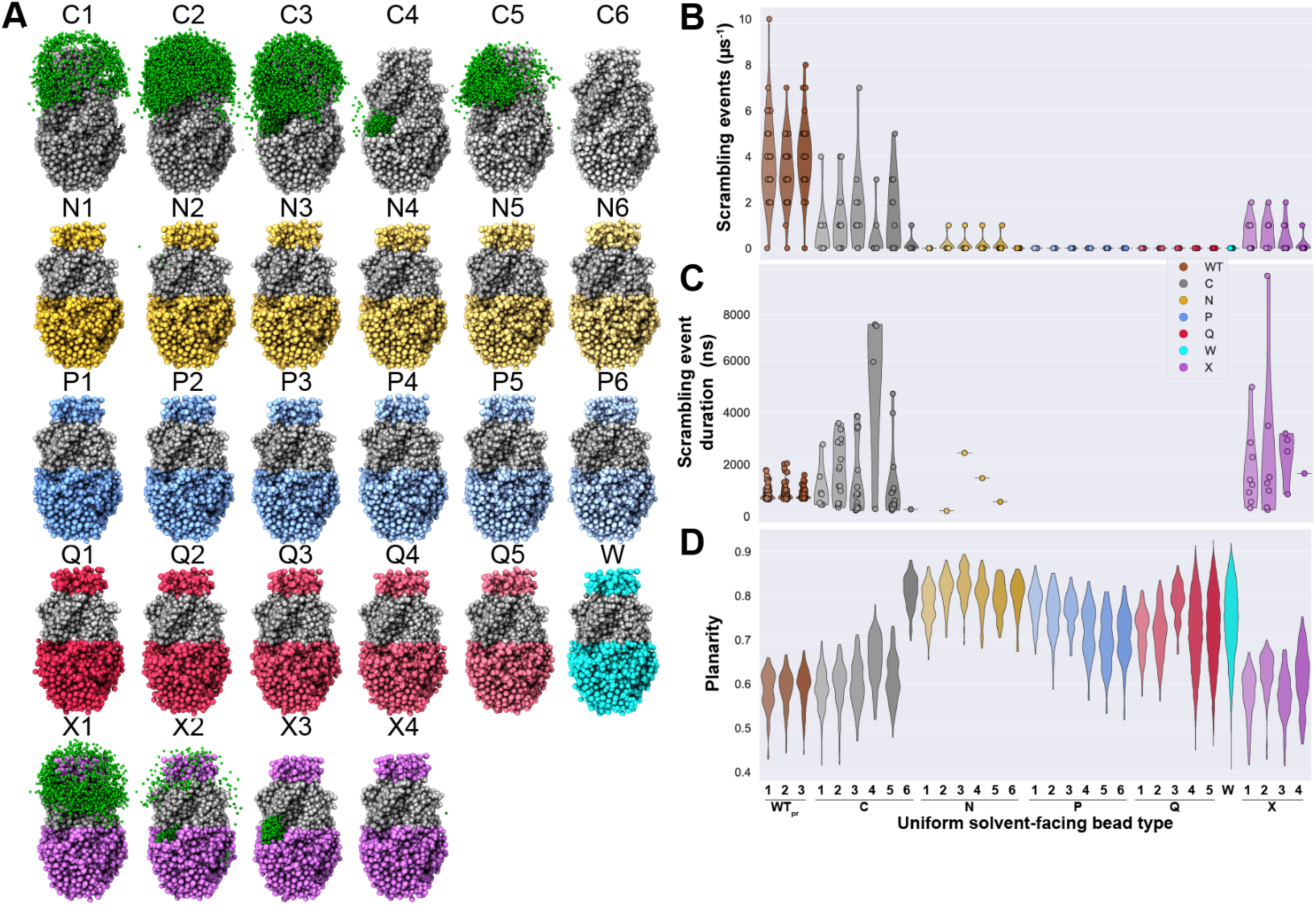
Scrambling by two-region systems with shape based on dimeric nhTMEM16. (*A*) The TM region of each nhTMEM16-based scramblase is modeled as bead type C4 (gray) while the solvent-facing beads vary (colors). Systems have been minimized and equilibrated. Green spheres illustrate the path taken by the head group during scrambling. (*B*–*D*) Scrambling rate, duration per event, and planarity for the systems shown in (*A*). Data for WTpr appears in Figures 4, 5, S5-8, and S10.

**Figure S7.**
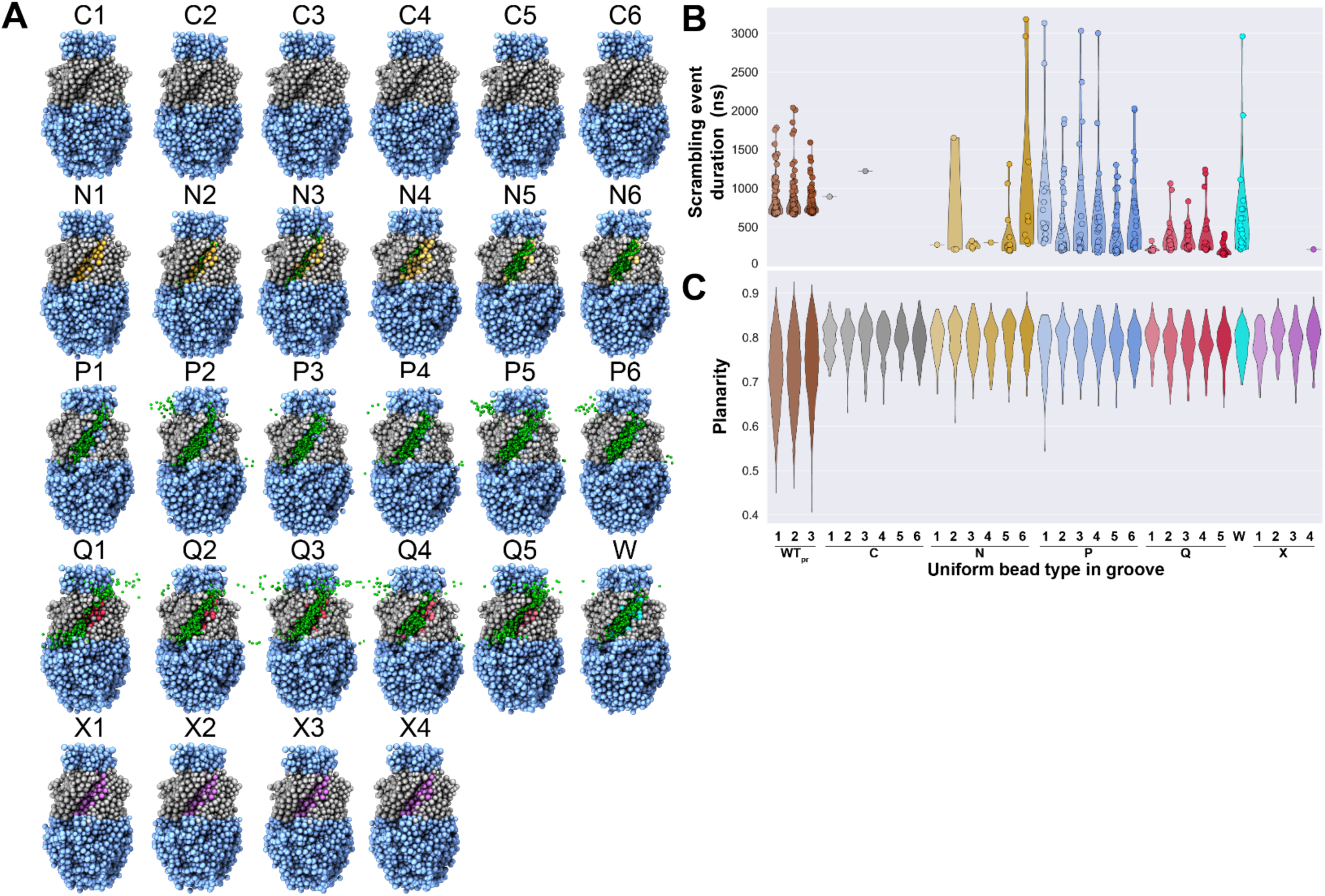
Scrambling by three-region systems with shape based on dimeric nhTMEM16. (*A*) The TM region of each nhTMEM16-based scramblase is modeled as bead type C4 (gray) while the solvent-facing beads are P2 (sky blue) and groove beads vary (colors). Systems have been minimized and equilibrated. Green spheres illustrate the path taken by the head group during scrambling. (*B, C*) Scrambling rate and planarity for the systems shown in (*A*). Models for C1, N1, and P2 appear in Figure 4. Data for WTpr appears in Figures 4, 5, S5-8, and S10.

**Figure S8.**
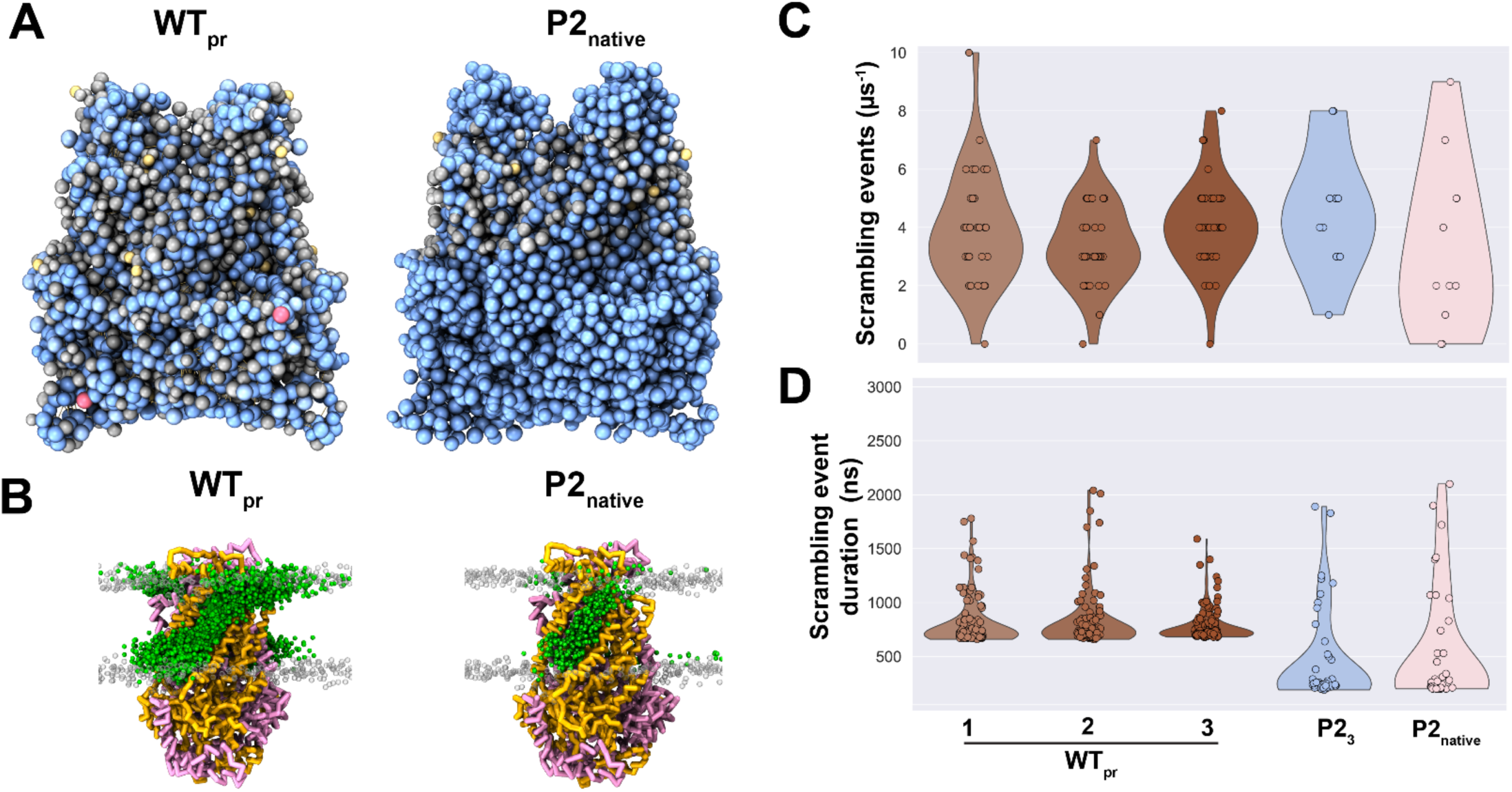
Bead types in the TMng region do not determine scrambling. (*A*, *B*) Systems are built with bead types completely determined by the native sequence (WTpr, *left*) or with P23-model beads everywhere except the TMng region (right). Transparent gray spheres mark phosphate head groups (bead PO4) and green spheres illustrate the path taken by the head groups during scrambling. (*C*, *D*) Rate and time per scrambling event for each system. Data for P23 appears in Figure 4I, 5B, and S10. Data for WTpr appears in Figures 4, 5, S5-8, and S10.

**Figure S9.**
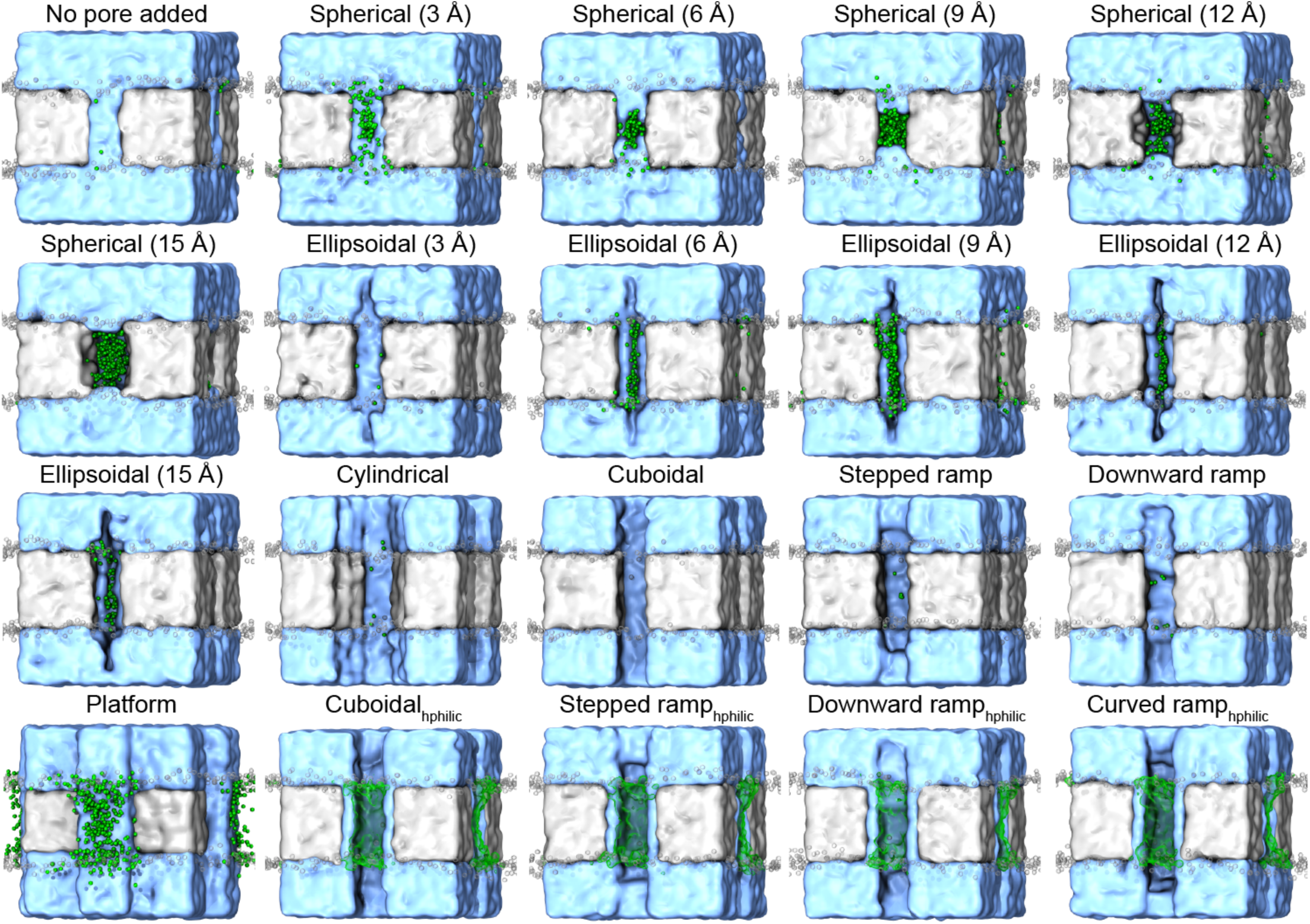
Groove architecture controls scrambling in simple cubic systems. Each cubic system is shown with hydrophilic beads (sky blue, type P2) and TMng beads (gray, type C4). Green spheres mark the paths taken by POPC head groups during scrambling events. Green densities are shown for POPC head groups for rapidly scrambling systems. We removed beads to form a groove with the indicated shape. For spherical grooves, the indicated length is the radius of the sphere. For ellipsoidal grooves the indicated length controls the depth of the groove toward the center of the cube. The cylindrical groove has radius 18 Å and the cuboidal groove has depth 25 Å. The downward ramp groove is most shallow at the bottom of the cube. The stepped and curved ramps are most shallow at the center of the cube. The platform extends outward from the box by ∼10 Å. Subscript *hphilic* indicates that the walls of the groove are comprised of hydrophilic (sky blue) beads. Some models shown here appear in Figure 7.

**Figure S10.**
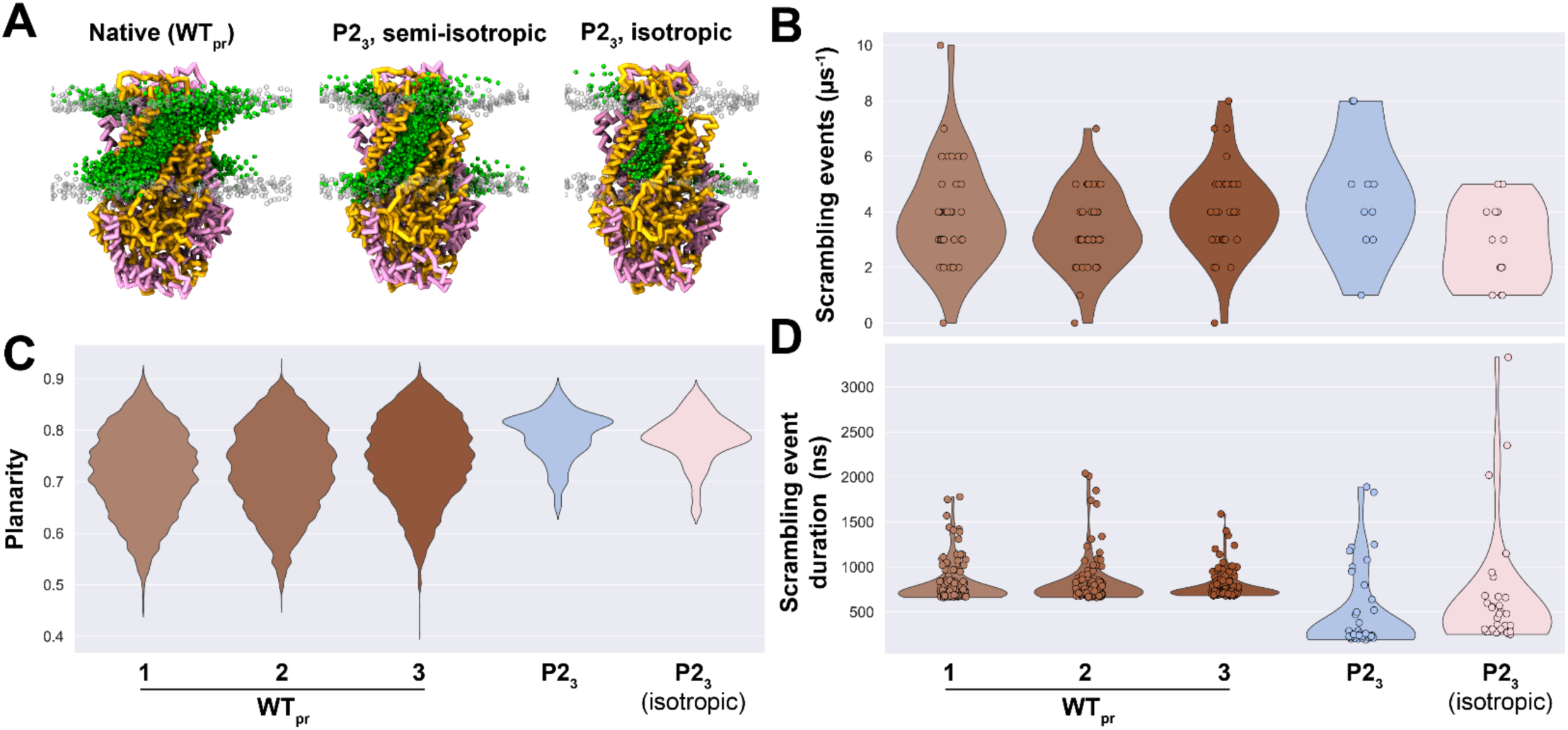
An isotropic barostat does not affect scrambling in simple Martini3 scramblases. (*A*) Scrambling pathways for WTpr (*left*) and the P23 model with a semiisotropic (*middle*) or isotropic barostat (*right*). (*B*–*D*) Planarity of the lipid bilayer alongside scrambling rate and time per event for each system. Data for P23 appears in Figure 4I, 5B, and S8. Data for WTpr appears in Figures 4, 5, S5-8, and S10.

**Figure S11.**
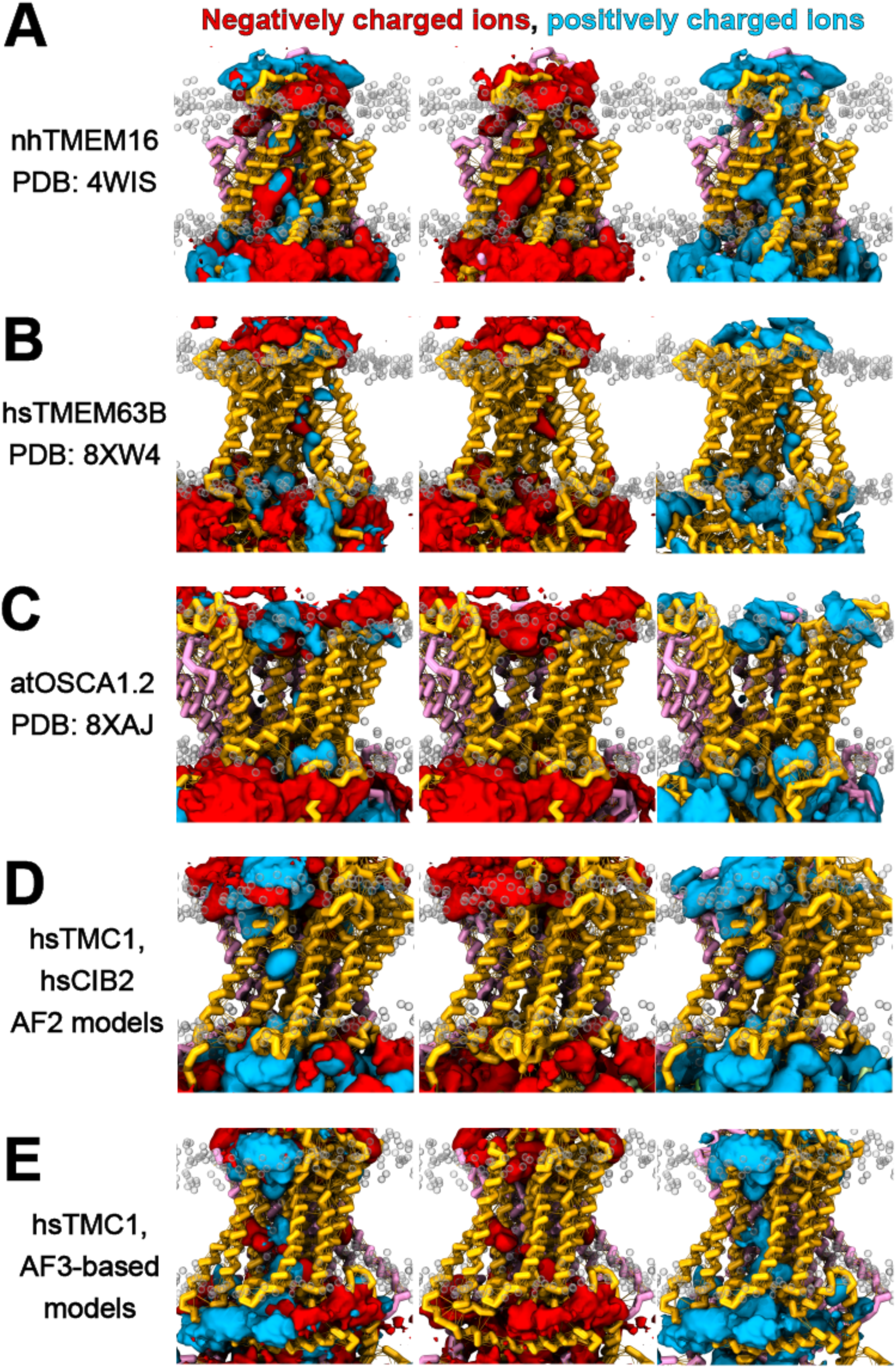
Ion occupancy for TCS family members using CG models. (*A*–*E*) Initial structures overlaid with ion densities. Negatively (red) and positively charged (blue) densities are shown together (*left*) or separately (*middle*, *right*). Transparent gray spheres mark non-scrambling phosphate head groups. Time resolution for density computation was 100 ps.

**Table S1.** Simulation parameters for TCS family proteins.

| Simulation | Time (μs) | $N_{\text{atoms}}$ | $x$ ($\text{\AA}$) | $y$ ($\text{\AA}$) | $z$ ($\text{\AA}$) | $N_{\text{w}}$ | $N_{\text{protein}}$ | $N_{\text{protres}}$ | $N_{\text{lipidres}}$ | $N_{\text{Cl}}$ | $N_{\text{Na}}$ | Lipid |
| --- | --- | --- | --- | --- | --- | --- | --- | --- | --- | --- | --- | --- |
| atOSCA1.2, dimer, closed | 10.5 | 65,402 | 183.2 | 183.2 | 220.5 | 50,190 | 3,522 | 1,434 | 882 | 565 | 541 | POPC |
| atOSCA1.2, dimer, open | 15.5 | 65,417 | 183.1 | 183.1 | 220.9 | 50,373 | 3,518 | 1,432 | 868 | 567 | 543 |  |
| atOSCA1.2, monomer, closed | 10.4 | 64,803 | 183.6 | 183.6 | 220.3 | 50,212 | 1,761 | 717 | 977 | 559 | 547 |  |
| hsTMC1 and hsCIB2, dimer | 10.1 | 63,888 | 181.2 | 181.2 | 218.2 | 47,906 | 4,308 | 1,774 | 885 | 534 | 520 |  |
| hsTMC1, dimer | 10.7 | 64,818 | 183.5 | 183.5 | 217.8 | 49,478 | 3,426 | 1,400 | 902 | 535 | 555 |  |
| hsTMC1, dimer, open | 12.2 | 64,920 | 183.4 | 183.4 | 218.5 | 49,804 | 3,316 | 1,354 | 892 | 553 | 543 |  |
| hsTMEM63A, closed | 10.1 | 64,955 | 183.8 | 183.8 | 220.2 | 50,250 | 1,718 | 700 | 990 | 555 | 552 |  |
| hsTMEM63B, open | 10.4 | 64,728 | 183.7 | 183.7 | 219.6 | 50,146 | 1,790 | 723 | 974 | 560 | 544 |  |
| hsTMEM63B, closed | 10.5 | 65,032 | 184.3 | 184.3 | 219.0 | 50,179 | 1,785 | 722 | 997 | 560 | 544 |  |
| hsTMEM63C, closed | 10.7 | 64,903 | 183.7 | 183.7 | 219.9 | 50,264 | 1,760 | 705 | 981 | 558 | 549 |  |
| nhTMEM16, dimer, closed | 8.7 | 65,527 | 183.9 | 183.9 | 219.0 | 49,737 | 3,462 | 1,442 | 936 | 538 | 558 |  |
| nhTMEM16, dimer, open | 12.5 | 65,462 | 183.8 | 183.8 | 219.3 | 49,826 | 3,462 | 1,442 | 923 | 539 | 559 |  |

**Table S2.** Simulation parameters for simple scramblases with the shape of nhTMEM16 and a uniform bead type for protein.

| Simulation | Time (μs) | $N_{\text{atoms}}$ | $x$ ($\text{\AA}$) | $y$ ($\text{\AA}$) | $z$ ($\text{\AA}$) | $N_w$ | $N_{\text{protein}}$ | $N_{\text{protres}}$ | $N_{\text{lipidres}}$ | $N_{\text{Cl}}$ | $N_{\text{Na}}$ | Lipid |
| --- | --- | --- | --- | --- | --- | --- | --- | --- | --- | --- | --- | --- |
| C1 | 10.7 | 65,476 | 183.1 | 183.1 | 224.3 | 49,818 | 3,462 | 1,442 | 925 | 548 | 548 | POPC |
| C2 | 11.4 | 65,465 | 182.4 | 182.4 | 225.6 | 49,807 |  |  |  |  |  |  |
| C3 | 11.7 | 65,454 | 182.5 | 182.5 | 225.0 | 49,796 |  |  |  |  |  |  |
| C4 | 9.9 | 65,461 | 182.5 | 182.5 | 225.1 | 49,803 |  |  |  |  |  |  |
| C5 | 12.0 | 65,455 | 182.8 | 182.8 | 224.2 | 49,797 |  |  |  |  |  |  |
| C6 | 11.2 | 65,478 | 182.6 | 182.6 | 224.5 | 49,820 |  |  |  |  |  |  |
| N1 | 10.9 | 65,458 | 183.1 | 183.1 | 223.2 | 49,800 |  |  |  |  |  |  |
| N2 | 11.0 | 65,458 | 182.8 | 182.8 | 223.3 | 49,800 |  |  |  |  |  |  |
| N3 | 11.3 | 65,455 | 182.4 | 182.4 | 224.4 | 49,797 |  |  |  |  |  |  |
| N4 | 9.9 | 65,464 | 182.7 | 182.7 | 223.7 | 49,806 |  |  |  |  |  |  |
| N5 | 10.8 | 65,472 | 182.3 | 182.3 | 224.4 | 49,814 |  |  |  |  |  |  |
| N6 | 11.4 | 65,468 | 182.7 | 182.7 | 223.6 | 49,810 |  |  |  |  |  |  |
| P1 | 12.0 | 65,448 | 182.5 | 182.5 | 223.9 | 49,790 |  |  |  |  |  |  |
| P2 | 9.4 | 65,470 | 182.7 | 182.7 | 223.5 | 49,812 |  |  |  |  |  |  |
| P3 | 12.0 | 65,446 | 182.4 | 182.4 | 224.0 | 49,788 |  |  |  |  |  |  |
| P4 | 9.8 | 65,453 | 183.1 | 183.1 | 222.1 | 49,795 |  |  |  |  |  |  |
| P5 | 10.5 | 65,453 | 182.9 | 182.9 | 222.6 | 49,795 |  |  |  |  |  |  |
| P6 | 6.7 | 65,467 | 182.7 | 182.7 | 222.8 | 49,809 |  |  |  |  |  |  |
| Q1 | 12.8 | 65,475 | 182.2 | 182.2 | 224.2 | 49,817 |  |  |  |  |  |  |
| Q2 | 12.3 | 65,457 | 182.4 | 182.4 | 223.6 | 49,799 |  |  |  |  |  |  |
| Q3 | 12.2 | 65,441 | 182.3 | 182.3 | 223.5 | 49,783 |  |  |  |  |  |  |
| Q4 | 12.5 | 65,462 | 182.4 | 182.4 | 223.1 | 49,804 |  |  |  |  |  |  |
| Q5 | 11.4 | 65,451 | 182.6 | 182.6 | 222.6 | 49,793 |  |  |  |  |  |  |
| W | 7.3 | 65,463 | 182.7 | 182.7 | 223.9 | 49,805 |  |  |  |  |  |  |
| X1 | 11.7 | 65,462 | 183.4 | 183.4 | 223.1 | 49,804 |  |  |  |  |  |  |
| X2 | 10.9 | 65,458 | 182.6 | 182.6 | 224.7 | 49,800 |  |  |  |  |  |  |
| X3 | 11.9 | 65,470 | 182.8 | 182.8 | 224.3 | 49,812 |  |  |  |  |  |  |
| X4 | 11.9 | 65,465 | 182.7 | 182.7 | 224.1 | 49,807 |  |  |  |  |  |  |

**Table S3.** Simulation parameters for nhTMEM16 with bead types of all residues determined by the native sequence and with positional restraints (WTpr).

| Simulation | Time (μs) | *N*<sub>atoms</sub> | *x* (Å) | *y* (Å) | *z* (Å) | *N*<sub>w</sub> | *N*<sub>protein</sub> | *N*<sub>protres</sub> | *N*<sub>lipidres</sub> | *N*<sub>Cl</sub> | *N*<sub>Na</sub> | Lipid |
| --- | --- | --- | --- | --- | --- | --- | --- | --- | --- | --- | --- | --- |
| WT 1 | 34.6 | 65,465 | 183.4 | 183.4 | 221.6 | 49,807 | 3,462 | 1,442 | 925 | 548 | 548 | POPC |
| WT 2 | 35.0 | 65,471 | 184.2 | 184.2 | 220.1 | 49,813 |  |  |  |  |  |  |
| WT 3 | 35.6 | 65,478 | 184.2 | 184.2 | 219.8 | 49,820 |  |  |  |  |  |  |

**Table S4.** Simulation parameters for simple scramblases with the shape of nhTMEM16 and two distinct regions. Bead type C4 is placed in the TM region, and the solvent-facing regions have a varying uniform bead type.

| Simulation | Time (μs) | $N_{\text{atoms}}$ | $x$ ($\text{\AA}$) | $y$ ($\text{\AA}$) | $z$ ($\text{\AA}$) | $N_w$ | $N_{\text{protein}}$ | $N_{\text{protres}}$ | $N_{\text{lipidres}}$ | $N_{\text{Cl}}$ | $N_{\text{Na}}$ | Lipid |
| --- | --- | --- | --- | --- | --- | --- | --- | --- | --- | --- | --- | --- |
| C1 | 12.3 | 65,468 | 184.3 | 184.3 | 221.6 | 49,810 | 3,462 | 1,442 | 925 | 548 | 548 | POPC |
| C2 | 11.3 | 65,446 | 184.5 | 184.5 | 220.8 | 49,788 |  |  |  |  |  |  |
| C3 | 10.7 | 65,466 | 184.0 | 184.0 | 221.5 | 49,808 |  |  |  |  |  |  |
| C4 | 11.2 | 65,471 | 184.3 | 184.3 | 221.1 | 49,813 |  |  |  |  |  |  |
| C5 | 11.5 | 65,452 | 184.1 | 184.1 | 221.6 | 49,794 |  |  |  |  |  |  |
| C6 | 9.7 | 65,468 | 184.2 | 184.2 | 221.1 | 49,810 |  |  |  |  |  |  |
| N1 | 11.4 | 65,456 | 184.0 | 184.0 | 220.9 | 49,798 |  |  |  |  |  |  |
| N2 | 10.9 | 65,466 | 183.7 | 183.7 | 221.6 | 49,808 |  |  |  |  |  |  |
| N3 | 11.4 | 65,462 | 183.7 | 183.7 | 221.6 | 49,804 |  |  |  |  |  |  |
| N4 | 11.1 | 65,446 | 183.7 | 183.7 | 221.3 | 49,788 |  |  |  |  |  |  |
| N5 | 10.3 | 65,467 | 183.5 | 183.5 | 221.7 | 49,809 |  |  |  |  |  |  |
| N6 | 11.1 | 65,463 | 183.7 | 183.7 | 221.1 | 49,805 |  |  |  |  |  |  |
| P1 | 11.3 | 65,463 | 184.1 | 184.1 | 220.2 | 49,805 |  |  |  |  |  |  |
| P2 | 11.9 | 65,463 | 183.7 | 183.7 | 221.1 | 49,805 |  |  |  |  |  |  |
| P3 | 11.0 | 65,455 | 183.7 | 183.7 | 221.1 | 49,797 |  |  |  |  |  |  |
| P4 | 8.2 | 65,450 | 183.8 | 183.8 | 220.8 | 49,792 |  |  |  |  |  |  |
| P5 | 10.5 | 65,459 | 183.0 | 183.0 | 222.4 | 49,801 |  |  |  |  |  |  |
| P6 | 10.3 | 65,476 | 183.2 | 183.2 | 222.1 | 49,818 |  |  |  |  |  |  |
| Q1 | 11.4 | 65,440 | 183.8 | 183.8 | 220.1 | 49,782 |  |  |  |  |  |  |
| Q2 | 10.8 | 65,452 | 183.3 | 183.3 | 221.5 | 49,794 |  |  |  |  |  |  |
| Q3 | 11.0 | 65,451 | 183.5 | 183.5 | 220.9 | 49,793 |  |  |  |  |  |  |
| Q4 | 11.5 | 65,455 | 183.5 | 183.5 | 220.6 | 49,797 |  |  |  |  |  |  |
| Q5 | 11.9 | 65,447 | 184.2 | 184.2 | 219.0 | 49,789 |  |  |  |  |  |  |
| W | 10.7 | 65,439 | 183.9 | 183.9 | 220.8 | 49,781 |  |  |  |  |  |  |
| X1 | 10.7 | 65,459 | 184.3 | 184.3 | 221.1 | 49,801 |  |  |  |  |  |  |
| X2 | 10.9 | 65,453 | 183.8 | 183.8 | 222.2 | 49,795 |  |  |  |  |  |  |
| X3 | 10.1 | 65,467 | 183.9 | 183.9 | 221.9 | 49,809 |  |  |  |  |  |  |
| X4 | 10.7 | 65,451 | 183.8 | 183.8 | 221.9 | 49,793 |  |  |  |  |  |  |

**Table S5.** Simulation parameters for simple scramblases with the shape of nhTMEM16 and three distinct regions. Bead type C4 is placed in the TMng region, the solvent-facing regions are comprised of bead P2, and the groove is built from a varying uniform bead type.

| Simulation | Time (μs) | $N_{\text{atoms}}$ | $x$ (Å) | $y$ (Å) | $z$ (Å) | $N_w$ | $N_{\text{protein}}$ | $N_{\text{protres}}$ | $N_{\text{lipidres}}$ | $N_{\text{Cl}}$ | $N_{\text{Na}}$ | Lipid |
| --- | --- | --- | --- | --- | --- | --- | --- | --- | --- | --- | --- | --- |
| C1 | 10.6 | 65,451 | 183.7 | 183.7 | 221.3 | 49,793 | 3,462 | 1,442 | 925 | 548 | 548 | POPC |
| C2 | 11.1 | 65,462 | 183.8 | 183.8 | 220.9 | 49,804 |  |  |  |  |  |  |
| C3 | 10.9 | 65,445 | 183.4 | 183.4 | 221.6 | 49,787 |  |  |  |  |  |  |
| C4 | 11.5 | 65,475 | 183.9 | 183.9 | 220.8 | 49,817 |  |  |  |  |  |  |
| C5 | 11.7 | 65,453 | 183.6 | 183.6 | 221.1 | 49,795 |  |  |  |  |  |  |
| C6 | 11.1 | 65,458 | 183.6 | 183.6 | 221.1 | 49,800 |  |  |  |  |  |  |
| N1 | 10.8 | 65,464 | 183.5 | 183.5 | 221.4 | 49,806 |  |  |  |  |  |  |
| N2 | 10.7 | 65,456 | 183.4 | 183.4 | 222.0 | 49,798 |  |  |  |  |  |  |
| N3 | 11.4 | 65,460 | 183.6 | 183.6 | 221.3 | 49,802 |  |  |  |  |  |  |
| N4 | 11.3 | 65,452 | 183.8 | 183.8 | 220.7 | 49,794 |  |  |  |  |  |  |
| N5 | 10.8 | 65,461 | 183.7 | 183.7 | 221.0 | 49,803 |  |  |  |  |  |  |
| N6 | 11.9 | 65,459 | 183.7 | 183.7 | 221.0 | 49,801 |  |  |  |  |  |  |
| P1 | 11.2 | 65,452 | 183.6 | 183.6 | 221.3 | 49,794 |  |  |  |  |  |  |
| P2 (P2<sub>3</sub>) | 10.8 | 65,460 | 183.6 | 183.6 | 221.4 | 49,802 |  |  |  |  |  |  |
| P3 | 9.8 | 65,452 | 183.5 | 183.5 | 221.6 | 49,794 |  |  |  |  |  |  |
| P4 | 9.8 | 65,466 | 183.8 | 183.8 | 221.0 | 49,808 |  |  |  |  |  |  |
| P5 | 9.0 | 65,462 | 183.3 | 183.3 | 221.9 | 49,804 |  |  |  |  |  |  |
| P6 | 11.4 | 65,450 | 183.9 | 183.9 | 220.8 | 49,792 |  |  |  |  |  |  |
| Q1 | 10.7 | 65,462 | 183.5 | 183.5 | 221.5 | 49,804 |  |  |  |  |  |  |
| Q2 | 11.3 | 65,446 | 183.5 | 183.5 | 221.3 | 49,788 |  |  |  |  |  |  |
| Q3 | 11.5 | 65,449 | 183.7 | 183.7 | 221.2 | 49,791 |  |  |  |  |  |  |
| Q4 | 11.5 | 65,464 | 183.6 | 183.6 | 221.0 | 49,806 |  |  |  |  |  |  |
| Q5 | 7.8 | 65,449 | 183.5 | 183.5 | 221.4 | 49,791 |  |  |  |  |  |  |
| W | 11.3 | 65,465 | 183.5 | 183.5 | 221.6 | 49,807 |  |  |  |  |  |  |
| X1 | 10.3 | 65,459 | 183.6 | 183.6 | 221.3 | 49,801 |  |  |  |  |  |  |
| X2 | 11.3 | 65,455 | 183.9 | 183.9 | 220.8 | 49,797 |  |  |  |  |  |  |
| X3 | 10.2 | 65,456 | 183.8 | 183.8 | 220.6 | 49,798 |  |  |  |  |  |  |
| X4 | 11.3 | 65,459 | 184.0 | 184.0 | 220.5 | 49,801 |  |  |  |  |  |  |

**Table S6.**
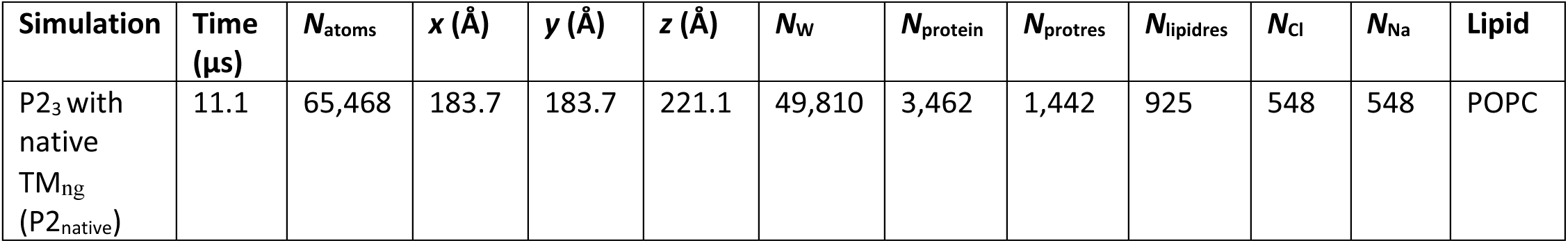
Simulation parameters for P23 with TMng beads determined by the native sequence of nhTMEM16 (P2native).

| Simulation | Time<br>(μs) | $N_{\text{atoms}}$ | $x$ (Å) | $y$ (Å) | $z$ (Å) | $N_{\text{w}}$ | $N_{\text{protein}}$ | $N_{\text{protres}}$ | $N_{\text{lipidres}}$ | $N_{\text{Cl}}$ | $N_{\text{Na}}$ | Lipid |
| --- | --- | --- | --- | --- | --- | --- | --- | --- | --- | --- | --- | --- |
| P2<sub>3</sub> with<br>native<br>TM<sub>ng</sub><br>(P2<sub>native</sub>) | 11.1 | 65,468 | 183.7 | 183.7 | 221.1 | 49,810 | 3,462 | 1,442 | 925 | 548 | 548 | POPC |

**Table S7.** Simulation parameters for the P2polyala system.

| Simulation | Time<br>(μs) | N<sub>atoms</sub> | x (Å) | y (Å) | z (Å) | N<sub>w</sub> | N<sub>protein</sub> | N<sub>protres</sub> | N<sub>lipidres</sub> | N<sub>Cl</sub> | N<sub>Na</sub> | Lipid |
| --- | --- | --- | --- | --- | --- | --- | --- | --- | --- | --- | --- | --- |
| P2<sub>polyala</sub> | 11.3 | 65,323 | 182.1 | 182.1 | 222.0 | 50,129 | 2,882 | 1,442 | 934 | 552 | 552 | POPC |

**Table S8.** Simulation parameters for P23 systems after rotating the hydrophilic groove and projecting it onto other parts of the protein surface.

| Simulation | Time<br>(μs) | $N_{\text{atoms}}$ | $x$ (Å) | $y$ (Å) | $z$ (Å) | $N_w$ | $N_{\text{protein}}$ | $N_{\text{protres}}$ | $N_{\text{lipidres}}$ | $N_{\text{Cl}}$ | $N_{\text{Na}}$ | Lipid |
| --- | --- | --- | --- | --- | --- | --- | --- | --- | --- | --- | --- | --- |
| 0 | 10.5 | 65,449 | 183.8 | 183.8 | 220.9 | 49,791 | 3,462 | 1,442 | 925 | 548 | 548 | POPC |
| 20 | 10.0 | 65,455 | 183.3 | 183.3 | 221.9 | 49,797 |  |  |  |  |  |  |
| 40 | 10.1 | 65,456 | 183.8 | 183.8 | 220.6 | 49,798 |  |  |  |  |  |  |
| 60 | 10.0 | 65,449 | 183.4 | 183.4 | 221.7 | 49,791 |  |  |  |  |  |  |
| 80 | 10.8 | 65,473 | 183.7 | 183.7 | 221.2 | 49,815 |  |  |  |  |  |  |
| 100 | 10.0 | 65,466 | 183.8 | 183.8 | 221.0 | 49,808 |  |  |  |  |  |  |
| 120 | 10.3 | 65,457 | 184.1 | 184.1 | 220.0 | 49,799 |  |  |  |  |  |  |
| 140 | 10.4 | 65,469 | 183.6 | 183.6 | 221.3 | 49,811 |  |  |  |  |  |  |
| 160 | 10.1 | 65,481 | 183.5 | 183.5 | 221.5 | 49,823 |  |  |  |  |  |  |

**Table S9.** Simulation parameters for P23-based systems with varied thickness of the TMng region.

| Thickness (Å) | Time (μs) | *N*<sub>atoms</sub> | *x* (Å) | *y* (Å) | *z* (Å) | *N*<sub>w</sub> | *N*<sub>protein</sub> | *N*<sub>protres</sub> | *N*<sub>lipidres</sub> | *N*<sub>Cl</sub> | *N*<sub>Na</sub> | Lipid |
| --- | --- | --- | --- | --- | --- | --- | --- | --- | --- | --- | --- | --- |
| 10 | 11.4 | 65,453 | 183.8 | 183.8 | 220.8 | 49,795 | 3,462 | 1,442 | 925 | 548 | 548 | POPC |
| 14 | 11.6 | 65,447 | 183.5 | 183.5 | 221.4 | 49,789 |  |  |  |  |  |  |
| 18 | 11.5 | 65,449 | 183.6 | 183.6 | 220.9 | 49,791 |  |  |  |  |  |  |
| 22 | 14.0 | 65,461 | 183.6 | 183.6 | 221.0 | 49,803 |  |  |  |  |  |  |
| 26 | 15.6 | 65,453 | 183.2 | 183.2 | 222.2 | 49,795 |  |  |  |  |  |  |
| 30 | 12.4 | 65,451 | 183.4 | 183.4 | 221.9 | 49,793 |  |  |  |  |  |  |
| 34 | 12.2 | 65,458 | 183.6 | 183.6 | 221.3 | 49,800 |  |  |  |  |  |  |
| 38 | 10.1 | 65,449 | 183.6 | 183.6 | 221.2 | 49,791 |  |  |  |  |  |  |
| 42 | 10.1 | 65,459 | 183.8 | 183.8 | 221.0 | 49,801 |  |  |  |  |  |  |
| 46 | 10.1 | 65,449 | 183.6 | 183.6 | 221.4 | 49,791 |  |  |  |  |  |  |
| 50 | 10.3 | 65,463 | 183.7 | 183.7 | 221.2 | 49,805 |  |  |  |  |  |  |
| 54 | 10.3 | 65,454 | 183.7 | 183.7 | 221.5 | 49,796 |  |  |  |  |  |  |

**Table S10.** Simulation parameters for P23-based systems with an artificial hydrophobic gate inserted in the TMng region.

| Simulation | Time (μs) | $N_{\text{atoms}}$ | $x$ (Å) | $y$ (Å) | $z$ (Å) | $N_{\text{w}}$ | $N_{\text{protein}}$ | $N_{\text{protres}}$ | $N_{\text{lipidres}}$ | $N_{\text{Cl}}$ | $N_{\text{Na}}$ | Lipid |
| --- | --- | --- | --- | --- | --- | --- | --- | --- | --- | --- | --- | --- |
| C1 | 10.1 | 65,470 | 183.4 | 183.4 | 221.9 | 49,812 | 3,462 | 1,442 | 925 | 548 | 548 | POPC |
| C2 | 12.1 | 65,453 | 183.4 | 183.4 | 221.4 | 49,795 |  |  |  |  |  |  |
| C3 | 10.2 | 65,450 | 183.7 | 183.7 | 221.0 | 49,792 |  |  |  |  |  |  |
| C4 | 10.2 | 65,450 | 183.7 | 183.7 | 221.1 | 49,792 |  |  |  |  |  |  |
| C5 | 10.3 | 65,482 | 183.8 | 183.8 | 220.9 | 49,822 |  |  |  | 549 | 549 |  |
| C6 | 10.9 | 65,451 | 183.8 | 183.8 | 220.8 | 49,793 |  |  |  | 548 | 548 |  |
| C1 (thin) | 10.3 | 65,465 | 183.7 | 183.7 | 221.1 | 49,807 |  |  |  |  |  |  |
| C2 (thin) | 11.4 | 65,460 | 183.5 | 183.5 | 221.3 | 49,802 |  |  |  |  |  |  |
| C3 (thin) | 10.0 | 65,454 | 183.4 | 183.4 | 221.7 | 49,796 |  |  |  |  |  |  |
| C4 (thin) | 9.8 | 65,462 | 183.5 | 183.5 | 221.4 | 49,804 |  |  |  |  |  |  |
| C5 (thin) | 10.4 | 65,458 | 183.5 | 183.5 | 221.6 | 49,800 |  |  |  |  |  |  |
| C6 (thin) | 10.1 | 65,460 | 183.6 | 183.6 | 221.4 | 49,802 |  |  |  |  |  |  |

**Table S11.**
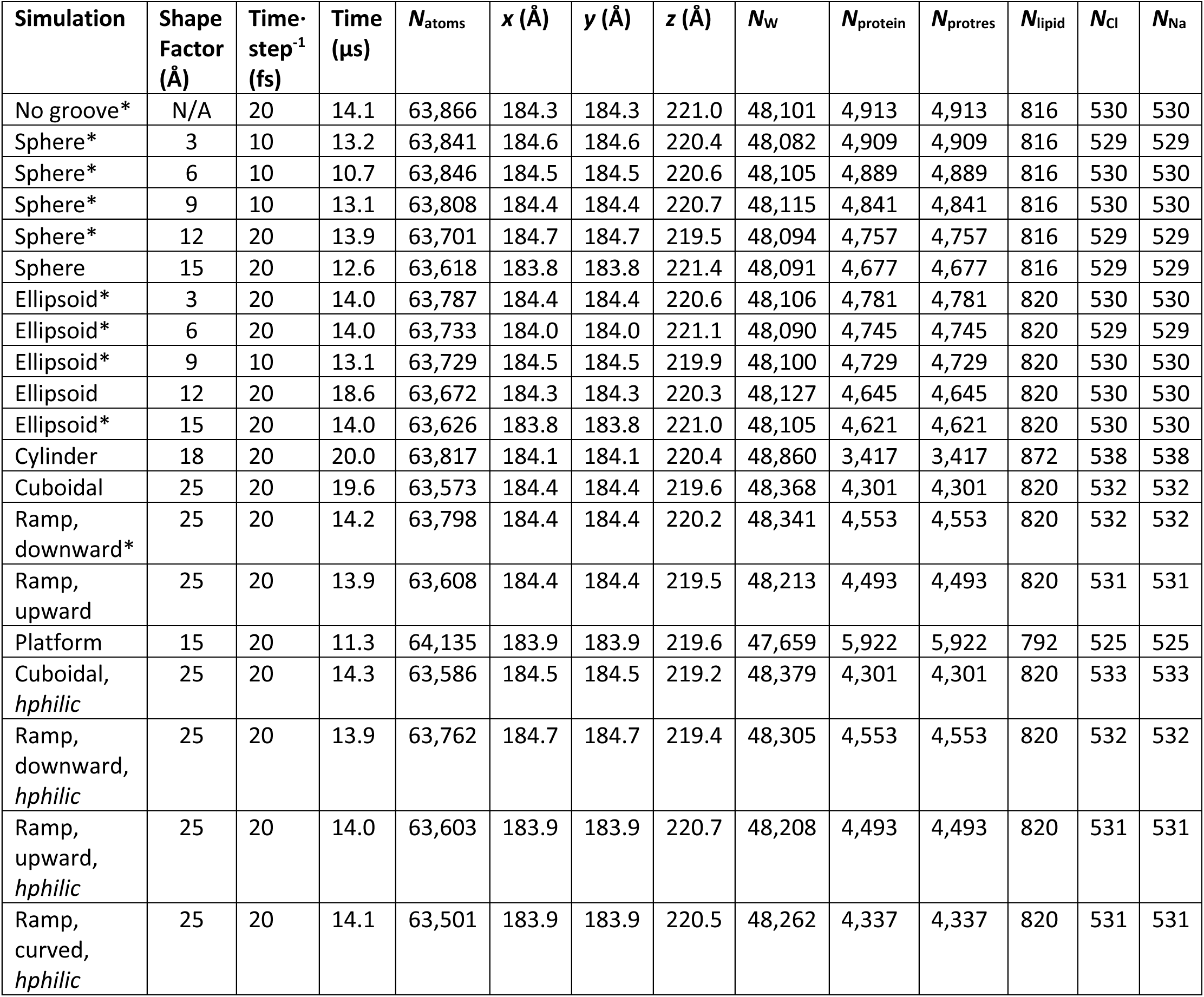
Simulation parameters for cubic systems with varied groove geometry. Simulation names with an asterisk (*) were run with an isotropic barostat. All systems contained POPC.

**Table S12.** Simulation parameters for P23 with pressure maintained isotropically.

| Simulation | Time<br>(μs) | $N_{\text{atoms}}$ | $x$ (Å) | $y$ (Å) | $z$ (Å) | $N_{\text{w}}$ | $N_{\text{protein}}$ | $N_{\text{protres}}$ | $N_{\text{lipidres}}$ | $N_{\text{Cl}}$ | $N_{\text{Na}}$ | Lipid |
| --- | --- | --- | --- | --- | --- | --- | --- | --- | --- | --- | --- | --- |
| P2<sub>3</sub> with<br>isotropic<br>barostat | 14.0 | 65,449 | 183.8 | 183.8 | 220.9 | 49,791 | 3,462 | 1,442 | 925 | 548 | 548 | POPC |

## REFERENCES

1. Devaux, P.F. 1991. Static and dynamic lipid asymmetry in cell membranes. Biochemistry. 30:1163–1173.

2. Ikeda, M., A. Kihara, and Y. Igarashi. 2006. Lipid Asymmetry of the Eukaryotic Plasma Membrane: Functions and Related Enzymes. Biol. Pharm. Bull. 29:1542–1546.

3. Lenoir, G., P. Williamson, and J.C. Holthuis. 2007. On the origin of lipid asymmetry: the flip side of ion transport. Curr. Opin. Chem. Biol. 11:654–661.

4. Sakuragi, T., and S. Nagata. 2023. Regulation of phospholipid distribution in the lipid bilayer by flippases and scramblases. Nat. Rev. Mol. Cell Biol. 24:576–596.

5. Sahu, S.K., S.N. Gummadi, N. Manoj, and G.K. Aradhyam. 2007. Phospholipid scramblases: An overview. Arch. Biochem. Biophys. 462:103–114.

6. Contreras, F.-X., L. Sánchez-Magraner, A. Alonso, and F.M. Goñi. 2010. Transbilayer (*flip-flop*) lipid motion and lipid scrambling in membranes. FEBS Lett. 584:1779–1786.

7. Williamson, P. 2015. Phospholipid Scramblases. Lipid Insights. 8s1:LPI.S31785.

8. Hankins, H.M., R.D. Baldridge, P. Xu, and T.R. Graham. 2015. Role of Flippases, Scramblases and Transfer Proteins in Phosphatidylserine Subcellular Distribution. Traffic. 16:35–47.

9. Lentz, B.R. 2003. Exposure of platelet membrane phosphatidylserine regulates blood coagulation. Prog. Lipid Res. 42:423–438.

10. Segawa, K., and S. Nagata. 2015. An Apoptotic ‘Eat Me’ Signal: Phosphatidylserine Exposure. Trends Cell Biol. 25:639–650.

11. McConnell, H.M., and R.D. Kornberg. 1971. Inside-outside transitions of phospholipids in vesicle membranes. Biochemistry. 10:1111–1120.

12. Parsegian, A. 1969. Energy of an Ion crossing a Low Dielectric Membrane: Solutions to Four Relevant Electrostatic Problems. Nature. 221:844–846.

13. Falzone, M.E., Z. Feng, O.E. Alvarenga, Y. Pan, B. Lee, X. Cheng, E. Fortea, S. Scheuring, and A. Accardi. 2022. TMEM16 scramblases thin the membrane to enable lipid scrambling. Nat. Commun. 13:2604.

14. Allhusen, J.S., and J.C. Conboy. 2017. The Ins and Outs of Lipid Flip-Flop. Acc. Chem. Res. 50:58–65.

15. Anglin, T.C., M.P. Cooper, H. Li, K. Chandler, and J.C. Conboy. 2010. Free Energy and Entropy of Activation for Phospholipid Flip-Flop in Planar Supported Lipid Bilayers. J. Phys. Chem. B. 114:1903–1914.

16. Anglin, T.C., and J.C. Conboy. 2008. Lateral Pressure Dependence of the Phospholipid Transmembrane Diffusion Rate in Planar-Supported Lipid Bilayers. Biophys. J. 95:186–193.

17. Anglin, T.C., and J.C. Conboy. 2009. Kinetics and Thermodynamics of Flip-Flop in Binary Phospholipid Membranes Measured by Sum-Frequency Vibrational Spectroscopy. Biochemistry. 48:10220–10234.

18. Brown, K.L., and J.C. Conboy. 2015. Phosphatidylglycerol Flip-Flop Suppression due to Headgroup Charge Repulsion. J. Phys. Chem. B. 119:10252–10260.

19. Liu, J., K.L. Brown, and J.C. Conboy. 2013. The effect of cholesterol on the intrinsic rate of lipid flip–flop as measured by sum-frequency vibrational spectroscopy. Faraday Discuss. 161:45–61.

20. Bevers, E.M., P. Comfurius, D.W.C. Dekkers, and R.F.A. Zwaal. 1999. Lipid translocation across the plasma membrane of mammalian cells. Biochim. Biophys. Acta BBA - Mol. Cell Biol. Lipids. 1439:317–330.

21. Kodigepalli, K.M., K. Bowers, A. Sharp, and M. Nanjundan. 2015. Roles and regulation of phospholipid scramblases. FEBS Lett. 589:3–14.

22. Sebinelli, H.G., C. Syska, A. Čopič, and G. Lenoir. 2024. Established and emerging players in phospholipid scrambling: A structural perspective. Biochimie. S0300908424002189.

23. Maeda, S., H. Yamamoto, L.N. Kinch, C.M. Garza, S. Takahashi, C. Otomo, N.V. Grishin, S. Forli, N. Mizushima, and T. Otomo. 2020. Structure, lipid scrambling activity and role in autophagosome formation of ATG9A. Nat. Struct. Mol. Biol. 27:1194–1201.

24. Matoba, K., T. Kotani, A. Tsutsumi, T. Tsuji, T. Mori, D. Noshiro, Y. Sugita, N. Nomura, S. Iwata, Y. Ohsumi, T. Fujimoto, H. Nakatogawa, M. Kikkawa, and N.N. Noda. 2020. Atg9 is a lipid scramblase that mediates autophagosomal membrane expansion. Nat. Struct. Mol. Biol. 27:1185–1193.

25. Huang, D., B. Xu, L. Liu, L. Wu, Y. Zhu, A. Ghanbarpour, Y. Wang, F.-J. Chen, J. Lyu, Y. Hu, Y. Kang, W. Zhou, X. Wang, W. Ding, X. Li, Z. Jiang, J. Chen, X. Zhang, H. Zhou, J.Z. Li, C. Guo, W. Zheng, X. Zhang, P. Li, T. Melia, K. Reinisch, and X.-W. Chen. 2021. TMEM41B acts as an ER scramblase required for lipoprotein biogenesis and lipid homeostasis. Cell Metab. 33:1655–1670.e8.

26. Li, Y.E., Y. Wang, X. Du, T. Zhang, H.Y. Mak, S.E. Hancock, H. McEwen, E. Pandzic, R.M. Whan, Y.C. Aw, I.E. Lukmantara, Y. Yuan, X. Dong, A. Don, N. Turner, S. Qi, and H. Yang. 2021. TMEM41B and VMP1 are scramblases and regulate the distribution of cholesterol and phosphatidylserine. J. Cell Biol. 220:e202103105.

27. Ma, Y., Y. Wang, X. Zhao, G. Jin, J. Xu, Z. Li, N. Yin, Z. Gao, B. Xia, and M. Peng. 2025. TMEM41B is an endoplasmic reticulum Ca2+ release channel maintaining naive T cell quiescence and responsiveness. Cell Discov. 11:18.

28. Morita, K., Y. Hama, and N. Mizushima. 2019. TMEM41B functions with VMP1 in autophagosome formation. Autophagy. 15:922–923.

29. Kalienkova, V., V. Clerico Mosina, and C. Paulino. 2021. The Groovy TMEM16 Family: Molecular Mechanisms of Lipid Scrambling and Ion Conduction. J. Mol. Biol. 433:166941.

30. Pedemonte, N., and L.J.V. Galietta. 2014. Structure and Function of TMEM16 Proteins (Anoctamins). Physiol. Rev. 94:419–459.

31. Bartoš, L., A.K. Menon, and R. Vácha. 2024. Insertases scramble lipids: Molecular simulations of MTCH2. Structure. 32:505–510.e4.

32. Li, D., C. Rocha-Roa, M.A. Schilling, K.M. Reinisch, and S. Vanni. 2024. Lipid scrambling is a general feature of protein insertases. Proc. Natl. Acad. Sci. 121:e2319476121.

33. Suzuki, J., E. Imanishi, and S. Nagata. 2014. Exposure of Phosphatidylserine by Xk-related Protein Family Members during Apoptosis. J. Biol. Chem. 289:30257–30267.

34. Jahn, H., L. Bartoš, G.I. Dearden, J.S. Dittman, J.C.M. Holthuis, R. Vácha, and A.K. Menon. 2023. Phospholipids are imported into mitochondria by VDAC, a dimeric beta barrel scramblase. Nat. Commun. 14:8115.

35. Rockenfeller, P. 2024. Phospholipid Scramblase Activity of VDAC Dimers: New Implications for Cell Death, Autophagy and Ageing. Biomolecules. 14:1218.

36. Goren, M.A., T. Morizumi, I. Menon, J.S. Joseph, J.S. Dittman, V. Cherezov, R.C. Stevens, O.P. Ernst, and A.K. Menon. 2014. Constitutive phospholipid scramblase activity of a G protein-coupled receptor. Nat. Commun. 5:5115.

37. Lowry, A.J., P. Liang, M. Song, Y. Wan, Z.-M. Pei, H. Yang, and Y. Zhang. 2024. TMEM16 and OSCA/TMEM63 proteins share a conserved potential to permeate ions and phospholipids. eLife. 13:RP96957.

38. Brunner, J.D., N.K. Lim, S. Schenck, A. Duerst, and R. Dutzler. 2014. X-ray structure of a calcium-activated TMEM16 lipid scramblase. Nature. 516:207–212.

39. Hahn. 2009. Anoctamin and transmembrane channel-like proteins are evolutionarily related. Int. J. Mol. Med. 24.

40. Jan, L.Y., and Y.N. Jan. 2025. Wide-ranging cellular functions of ion channels and lipid scramblases in the structurally related TMC, TMEM16 and TMEM63 families. Nat. Struct. Mol. Biol. 32:222–236.

41. Medrano-Soto, A., G. Moreno-Hagelsieb, D. McLaughlin, Z.S. Ye, K.J. Hendargo, and M.H. Saier. 2018. Bioinformatic characterization of the Anoctamin Superfamily of Ca2+-activated ion channels and lipid scramblases. PLOS ONE. 13:e0192851.

42. Kunzelmann, K., I. Cabrita, P. Wanitchakool, J. Ousingsawat, L. Sirianant, R. Benedetto, and R. Schreiber. 2016. Modulating Ca2+ signals: a common theme for TMEM16, Ist2, and TMC. Pflüg. Arch. - Eur. J. Physiol. 468:475–490.

43. Jeong, H., S. Clark, A. Goehring, S. Dehghani-Ghahnaviyeh, A. Rasouli, E. Tajkhorshid, and E. Gouaux. 2022. Structures of the TMC-1 complex illuminate mechanosensory transduction. Nature. 610:796–803.

44. Clark, S., H. Jeong, R. Posert, A. Goehring, and E. Gouaux. 2024. The structure of the *Caenorhabditis elegans* TMC-2 complex suggests roles of lipid-mediated subunit contacts in mechanosensory transduction. Proc. Natl. Acad. Sci. 121:e2314096121.

45. Pan, B., N. Akyuz, X.-P. Liu, Y. Asai, C. Nist-Lund, K. Kurima, B.H. Derfler, B. György, W. Limapichat, S. Walujkar, L.N. Wimalasena, M. Sotomayor, D.P. Corey, and J.R. Holt. 2018. TMC1 Forms the Pore of Mechanosensory Transduction Channels in Vertebrate Inner Ear Hair Cells. Neuron. 99:736–753.e6.

46. Giese, A.P., W.-H. Weng, K.S. Kindt, H.H.V. Chang, J.S. Montgomery, E.M. Ratzan, A.J. Beirl, R. Aponte Rivera, J.M. Lotthammer, S. Walujkar, M.P. Foster, O.A. Zobeiri, J.R. Holt, S. Riazuddin, K.E. Cullen, M. Sotomayor, and Z.M. Ahmed. 2025. Complexes of vertebrate TMC1/2 and CIB2/3 proteins form hair-cell mechanotransduction cation channels. eLife. 12:RP89719.

47. Ballesteros, A., and K.J. Swartz. 2018. Lipids surf the groove in scramblases. Proc. Natl. Acad. Sci. 115:7648–7650.

48. Ballesteros, A., C. Fenollar-Ferrer, and K.J. Swartz. 2018. Structural relationship between the putative hair cell mechanotransduction channel TMC1 and TMEM16 proteins. eLife. 7:e38433.

49. Ballesteros, A., and K.J. Swartz. 2022. Regulation of membrane homeostasis by TMC1 mechanoelectrical transduction channels is essential for hearing. Sci. Adv. 8:eabm5550.

50. Ebrahim, S., A. Ballesteros, W. Sharon Zheng, S. Mukherjee, G. Hu, W.-H. Weng, J.S. Montgomery, Y. Agyemang, R. Cui, W. Sun, E. Krystofiak, M.P. Foster, M. Sotomayor, and B. Kachar. 2024. Transmembrane channel-like 4 and 5 proteins at microvillar tips are potential ion channels and lipid scramblases. bioRxiv, doi: 10.1101/2024.08.22.609173 (preprint posted August 23, 2024).

51. Zheng, W., S. Rawson, Z. Shen, E. Tamilselvan, H.E. Smith, J. Halford, C. Shen, S.E. Murthy, M.H. Ulbrich, M. Sotomayor, T.-M. Fu, and J.R. Holt. 2023. TMEM63 proteins function as monomeric high-threshold mechanosensitive ion channels. Neuron. 111:3195–3210.e7.

52. Yuan, F., H. Yang, Y. Xue, D. Kong, R. Ye, C. Li, J. Zhang, L. Theprungsirikul, T. Shrift, B. Krichilsky, D.M. Johnson, G.B. Swift, Y. He, J.N. Siedow, and Z.-M. Pei. 2014. OSCA1 mediates osmotic-stress-evoked Ca2+ increases vital for osmosensing in Arabidopsis. Nature. 514:367–371.

53. Han, Y., Z. Zhou, R. Jin, F. Dai, Y. Ge, X. Ju, X. Ma, S. He, L. Yuan, Y. Wang, W. Yang, X. Yue, Z. Chen, Y. Sun, B. Corry, C.D. Cox, and Y. Zhang. 2024. Mechanical activation opens a lipid-lined pore in OSCA ion channels. Nature. 628:910–918.

54. Zhang, M., Y. Shan, C.D. Cox, and D. Pei. 2023. A mechanical-coupling mechanism in OSCA/TMEM63 channel mechanosensitivity. Nat. Commun. 14:3943.

55. Murthy, S.E., A.E. Dubin, T. Whitwam, S. Jojoa-Cruz, S.M. Cahalan, S.A.R. Mousavi, A.B. Ward, and A. Patapoutian. 2018. OSCA/TMEM63 are an evolutionarily conserved family of mechanically activated ion channels. eLife. 7:e41844.

56. Hou, C., W. Tian, T. Kleist, K. He, V. Garcia, F. Bai, Y. Hao, S. Luan, and L. Li. 2014. DUF221 proteins are a family of osmosensitive calcium-permeable cation channels conserved across eukaryotes. Cell Res. 24:632–635.

57. Zheng, W., A.J. Lowry, H.E. Smith, J. Xie, S. Rawson, C. Wang, J. Ou, M. Sotomayor, T.-M. Fu, H. Yang, and J.R. Holt. 2025. Structural and functional basis of mechanosensitive TMEM63 channelopathies. Neuron. S0896627325003563.

58. Wu, X., T. Shang, X. Lü, D. Luo, and D. Yang. 2024. A monomeric structure of human TMEM63A protein. Proteins Struct. Funct. Bioinforma. 92:750–756.

59. Miyata, Y., K. Takahashi, Y. Lee, C.S. Sultan, R. Kuribayashi, M. Takahashi, K. Hata, T. Bamba, Y. Izumi, K. Liu, T. Uemura, N. Nomura, S. Iwata, S. Nagata, T. Nishizawa, and K. Segawa. 2024. Membrane structure-responsive lipid scrambling by TMEM63B to control plasma membrane lipid distribution. Nat. Struct. Mol. Biol.

60. Jojoa-Cruz, S., K. Saotome, S.E. Murthy, C.C.A. Tsui, M.S. Sansom, A. Patapoutian, and A.B. Ward. 2018. Cryo-EM structure of the mechanically activated ion channel OSCA1.2. eLife. 7:e41845.

61. Jojoa-Cruz, S., A.E. Dubin, W.-H. Lee, and A.B. Ward. 2024. Structure-guided mutagenesis of OSCAs reveals differential activation to mechanical stimuli. eLife. 12:RP93147.

62. Qin, Y., D. Yu, D. Wu, J. Dong, W.T. Li, C. Ye, K.C. Cheung, Y. Zhang, Y. Xu, Y. Wang, Y.S. Shi, and S. Dang. 2023. Cryo-EM structure of TMEM63C suggests it functions as a monomer. Nat. Commun. 14:7265.

63. Jojoa-Cruz, S., B. Burendei, W.-H. Lee, and A.B. Ward. 2023. Structure of mechanically activated ion channel OSCA2.3 reveals mobile elements in the transmembrane domain. bioRxiv, doi: 10.1101/2023.06.15.545135 (preprint posted June 15, 2023).

64. Shan, Y., M. Zhang, M. Chen, X. Guo, Y. Li, M. Zhang, and D. Pei. 2024. Activation mechanisms of dimeric mechanosensitive OSCA/TMEM63 channels. Nat. Commun. 15:7504.

65. Zhang, M., D. Wang, Y. Kang, J.-X. Wu, F. Yao, C. Pan, Z. Yan, C. Song, and L. Chen. 2018. Structure of the mechanosensitive OSCA channels. Nat. Struct. Mol. Biol. 25:850–858.

66. Maity, K., J.M. Heumann, A.P. McGrath, N.J. Kopcho, P.-K. Hsu, C.-W. Lee, J.H. Mapes, D. Garza, S. Krishnan, G.P. Morgan, K.J. Hendargo, T. Klose, S.D. Rees, A. Medrano-Soto, M.H. Saier, M. Piñeros, E.A. Komives, J.I. Schroeder, G. Chang, and M.H.B. Stowell. 2019. Cryo-EM structure of OSCA1.2 from *Oryza sativa* elucidates the mechanical basis of potential membrane hyperosmolality gating. Proc. Natl. Acad. Sci. 116:14309–14318.

67. Liu, X., J. Wang, and L. Sun. 2018. Structure of the hyperosmolality-gated calcium-permeable channel OSCA1.2. Nat. Commun. 9:5060.

68. Paulino, C., V. Kalienkova, A.K.M. Lam, Y. Neldner, and R. Dutzler. 2017. Activation mechanism of the calcium-activated chloride channel TMEM16A revealed by cryo-EM. Nature. 552:421–425.

69. Feng, Z., O.E. Alvarenga, and A. Accardi. 2024. Structural basis of closed groove scrambling by a TMEM16 protein. Nat. Struct. Mol. Biol. 31:1468–1481.

70. Falzone, M.E., J. Rheinberger, B.-C. Lee, T. Peyear, L. Sasset, A.M. Raczkowski, E.T. Eng, A. Di Lorenzo, O.S. Andersen, C.M. Nimigean, and A. Accardi. 2019. Structural basis of Ca2+-dependent activation and lipid transport by a TMEM16 scramblase. eLife. 8:e43229.

71. Paulino, C., Y. Neldner, A.K. Lam, V. Kalienkova, J.D. Brunner, S. Schenck, and R. Dutzler. 2017. Structural basis for anion conduction in the calcium-activated chloride channel TMEM16A. eLife. 6:e26232.

72. Lam, A.K., and R. Dutzler. 2023. Mechanistic basis of ligand efficacy in the calcium-activated chloride channel TMEM16A. EMBO J. 42:e115030.

73. Arndt, M., C. Alvadia, M.S. Straub, V. Clerico Mosina, C. Paulino, and R. Dutzler. 2022. Structural basis for the activation of the lipid scramblase TMEM16F. Nat. Commun. 13:6692.

74. Lam, A.K.M., S. Rutz, and R. Dutzler. 2022. Inhibition mechanism of the chloride channel TMEM16A by the pore blocker 1PBC. Nat. Commun. 13:2798.

75. Dang, S., S. Feng, J. Tien, C.J. Peters, D. Bulkley, M. Lolicato, J. Zhao, K. Zuberbühler, W. Ye, L. Qi, T. Chen, C.S. Craik, Y.N. Jan, D.L. Minor, Y. Cheng, and L.Y. Jan. 2017. Cryo-EM structures of the TMEM16A calcium-activated chloride channel. Nature. 552:426–429.

76. Feng, Z., E. Di Zanni, O. Alvarenga, S. Chakraborty, N. Rychlik, and A. Accardi. 2024. In or out of the groove? Mechanisms of lipid scrambling by TMEM16 proteins. Cell Calcium. 121:102896.

77. Alvadia, C., N.K. Lim, V. Clerico Mosina, G.T. Oostergetel, R. Dutzler, and C. Paulino. 2019. Cryo-EM structures and functional characterization of the murine lipid scramblase TMEM16F. eLife. 8:e44365.

78. Kalienkova, V., V. Clerico Mosina, L. Bryner, G.T. Oostergetel, R. Dutzler, and C. Paulino. 2019. Stepwise activation mechanism of the scramblase nhTMEM16 revealed by cryo-EM. eLife. 8:e44364.

79. Bushell, S.R., A.C.W. Pike, M.E. Falzone, N.J.G. Rorsman, C.M. Ta, R.A. Corey, T.D. Newport, J.C. Christianson, L.F. Scofano, C.A. Shintre, A. Tessitore, A. Chu, Q. Wang, L. Shrestha, S.M.M. Mukhopadhyay, J.D. Love, N.A. Burgess-Brown, R. Sitsapesan, P.J. Stansfeld, J.T. Huiskonen, P. Tammaro, A. Accardi, and E.P. Carpenter. 2019. The structural basis of lipid scrambling and inactivation in the endoplasmic reticulum scramblase TMEM16K. Nat. Commun. 10:3956.

80. Lam, A.K.M., J. Rheinberger, C. Paulino, and R. Dutzler. 2021. Gating the pore of the calcium-activated chloride channel TMEM16A. Nat. Commun. 12:785.

81. Feng, S., C. Puchades, J. Ko, H. Wu, Y. Chen, E.E. Figueroa, S. Gu, T.W. Han, B. Ho, T. Cheng, J. Li, B. Shoichet, Y.N. Jan, Y. Cheng, and L.Y. Jan. 2023. Identification of a drug binding pocket in TMEM16F calcium-activated ion channel and lipid scramblase. Nat. Commun. 14:4874.

82. Feng, S., S. Dang, T.W. Han, W. Ye, P. Jin, T. Cheng, J. Li, Y.N. Jan, L.Y. Jan, and Y. Cheng. 2019. Cryo-EM Studies of TMEM16F Calcium-Activated Ion Channel Suggest Features Important for Lipid Scrambling. Cell Rep. 28:567–579.e4.

83. Giese, A.P.J., Y.-Q. Tang, G.P. Sinha, M.R. Bowl, A.C. Goldring, A. Parker, M.J. Freeman, S.D.M. Brown, S. Riazuddin, R. Fettiplace, W.R. Schafer, G.I. Frolenkov, and Z.M. Ahmed. 2017. CIB2 interacts with TMC1 and TMC2 and is essential for mechanotransduction in auditory hair cells. Nat. Commun. 8:43.

84. Pomorski, T., and A.K. Menon. 2006. Lipid flippases and their biological functions. Cell. Mol. Life Sci. 63:2908–2921.

85. Khelashvili, G., M.E. Falzone, X. Cheng, B.-C. Lee, A. Accardi, and H. Weinstein. 2019. Dynamic modulation of the lipid translocation groove generates a conductive ion channel in Ca2+-bound nhTMEM16. Nat. Commun. 10:4972.

86. Jia, Z., J. Huang, and J. Chen. 2022. Activation of TMEM16F by inner gate charged mutations and possible lipid/ion permeation mechanisms. Biophys. J. 121:3445–3457.

87. Le, T., Z. Jia, S.C. Le, Y. Zhang, J. Chen, and H. Yang. 2019. An inner activation gate controls TMEM16F phospholipid scrambling. Nat. Commun. 10:1846.

88. Bethel, N.P., and M. Grabe. 2016. Atomistic insight into lipid translocation by a TMEM16 scramblase. Proc. Natl. Acad. Sci. 113:14049–14054.

89. Lee, B.-C., G. Khelashvili, M. Falzone, A.K. Menon, H. Weinstein, and A. Accardi. 2018. Gating mechanism of the extracellular entry to the lipid pathway in a TMEM16 scramblase. Nat. Commun. 9:3251.

90. Kostritskii, A.Y., and J.-P. Machtens. 2021. Molecular mechanisms of ion conduction and ion selectivity in TMEM16 lipid scramblases. Nat. Commun. 12:2826.

91. Stephens, C.A., N. Van Hilten, L. Zheng, and M. Grabe. 2025. Simulation-based survey of TMEM16 family reveals that robust lipid scrambling requires an open groove. eLife, doi: 10.7554/eLife.105111.1 (preprint posted June 12, 2025).

92. Malvezzi, M., K.K. Andra, K. Pandey, B.-C. Lee, M.E. Falzone, A. Brown, R. Iqbal, A.K. Menon, and A. Accardi. 2018. Out-of-the-groove transport of lipids by TMEM16 and GPCR scramblases. Proc. Natl. Acad. Sci. 115.

93. Jiang, T., K. Yu, H.C. Hartzell, and E. Tajkhorshid. 2017. Lipids and ions traverse the membrane by the same physical pathway in the nhTMEM16 scramblase. eLife. 6:e28671.

94. Walujkar, S., J.M. Lotthammer, C.R. Nisler, J.C. Sudar, A. Ballesteros, and M. Sotomayor. 2021. *In Silico* Electrophysiology of Inner-Ear Mechanotransduction Channel TMC1 Models. bioRxiv, doi: 10.1101/2021.09.17.460860 *(preprint posted September 18, 2021)*.

95. Peineau, T., I. Marcovich, C.V.M. Rodriguez, S. O’Malley, R. Cui, A. Ballesteros, and J.R. Holt. 2025. Mammalian TMC1 or 2 are necessary for scramblase activity in auditory hair cells. Hear. Res. 460:109229.

96. Gyobu, S., K. Ishihara, J. Suzuki, K. Segawa, and S. Nagata. 2017. Characterization of the scrambling domain of the TMEM16 family. Proc. Natl. Acad. Sci. 114:6274–6279.

97. Niu, H., M. Maruoka, Y. Noguchi, H. Kosako, and J. Suzuki. 2024. Phospholipid scrambling induced by an ion channel/metabolite transporter complex. Nat. Commun. 15:7566.

98. Malvezzi, M., M. Chalat, R. Janjusevic, A. Picollo, H. Terashima, A.K. Menon, and A. Accardi. 2013. Ca2+-dependent phospholipid scrambling by a reconstituted TMEM16 ion channel. Nat. Commun. 4:2367.

99. Suzuki, J., M. Umeda, P.J. Sims, and S. Nagata. 2010. Calcium-dependent phospholipid scrambling by TMEM16F. Nature. 468:834–838.

100. Suzuki, J., T. Fujii, T. Imao, K. Ishihara, H. Kuba, and S. Nagata. 2013. Calcium-dependent Phospholipid Scramblase Activity of TMEM16 Protein Family Members. J. Biol. Chem. 288:13305–13316.

101. Gyobu, S., H. Miyata, M. Ikawa, D. Yamazaki, H. Takeshima, J. Suzuki, and S. Nagata. 2016. A Role of TMEM16E Carrying a Scrambling Domain in Sperm Motility. Mol. Cell. Biol. 36:645–659.

102. Yu, K., J.M. Whitlock, K. Lee, E.A. Ortlund, Y. Yuan Cui, and H.C. Hartzell. 2015. Identification of a lipid scrambling domain in ANO6/TMEM16F. eLife. 4:e06901.

103. Watanabe, R., T. Sakuragi, H. Noji, and S. Nagata. 2018. Single-molecule analysis of phospholipid scrambling by TMEM16F. Proc. Natl. Acad. Sci. 115:3066–3071.

104. Boccaccio, A., C. Picco, E. Di Zanni, and J. Scholz-Starke. 2022. Phospholipid scrambling by a TMEM16 homolog of *Arabidopsis thaliana*. FEBS J. 289:2578–2592.

105. Kim, J.-E., W. Ko, S. Jin, J.-N. Woo, Y. Jung, I. Bae, H.-K. Choe, D. Seo, B. Hille, and B.-C. Suh. 2025. Activation of TMEM16E scramblase induces ligand independent growth factor receptor signaling and macropinocytosis for membrane repair. Commun. Biol. 8:35.

106. Souza, P.C.T., R. Alessandri, J. Barnoud, S. Thallmair, I. Faustino, F. Grünewald, I. Patmanidis, H. Abdizadeh, B.M.H. Bruininks, T.A. Wassenaar, P.C. Kroon, J. Melcr, V. Nieto, V. Corradi, H.M. Khan, J. Domański, M. Javanainen, H. Martinez-Seara, N. Reuter, R.B. Best, I. Vattulainen, L. Monticelli, X. Periole, D.P. Tieleman, A.H. De Vries, and S.J. Marrink. 2021. Martini 3: a general purpose force field for coarse-grained molecular dynamics. Nat. Methods. 18:382–388.

107. Jumper, J., R. Evans, A. Pritzel, T. Green, M. Figurnov, O. Ronneberger, K. Tunyasuvunakool, R. Bates, A. Žídek, A. Potapenko, A. Bridgland, C. Meyer, S.A.A. Kohl, A.J. Ballard, A. Cowie, B. Romera-Paredes, S. Nikolov, R. Jain, J. Adler, T. Back, S. Petersen, D. Reiman, E. Clancy, M. Zielinski, M. Steinegger, M. Pacholska, T. Berghammer, S. Bodenstein, D. Silver, O. Vinyals, A.W. Senior, K. Kavukcuoglu, P. Kohli, and D. Hassabis. 2021. Highly accurate protein structure prediction with AlphaFold. Nature. 596:583–589.

108. Evans, R., M. O’Neill, A. Pritzel, N. Antropova, A. Senior, T. Green, A. Žídek, R. Bates, S. Blackwell, J. Yim, O. Ronneberger, S. Bodenstein, M. Zielinski, A. Bridgland, A. Potapenko, A. Cowie, K. Tunyasuvunakool, R. Jain, E. Clancy, P. Kohli, J. Jumper, and D. Hassabis. 2021. Protein complex prediction with AlphaFold-Multimer. bioRxiv, doi: 10.1101/2021.10.04.463034 *(preprint posted March 10, 2022)*.

109. Abramson, J., J. Adler, J. Dunger, R. Evans, T. Green, A. Pritzel, O. Ronneberger, L. Willmore, A.J. Ballard, J. Bambrick, S.W. Bodenstein, D.A. Evans, C.-C. Hung, M. O’Neill, D. Reiman, K. Tunyasuvunakool, Z. Wu, A. Žemgulytė, E. Arvaniti, C. Beattie, O. Bertolli, A. Bridgland, A. Cherepanov, M. Congreve, A.I. Cowen-Rivers, A. Cowie, M. Figurnov, F.B. Fuchs, H. Gladman, R. Jain, Y.A. Khan, C.M.R. Low, K. Perlin, A. Potapenko, P. Savy, S. Singh, A. Stecula, A. Thillaisundaram, C. Tong, S. Yakneen, E.D. Zhong, M. Zielinski, A. Žídek, V. Bapst, P. Kohli, M. Jaderberg, D. Hassabis, and J.M. Jumper. 2024. Accurate structure prediction of biomolecular interactions with AlphaFold 3. Nature. 630:493–500.

110. Varadi, M., S. Anyango, M. Deshpande, S. Nair, C. Natassia, G. Yordanova, D. Yuan, O. Stroe, G. Wood, A. Laydon, A. Žídek, T. Green, K. Tunyasuvunakool, S. Petersen, J. Jumper, E. Clancy, R. Green, A. Vora, M. Lutfi, M. Figurnov, A. Cowie, N. Hobbs, P. Kohli, G. Kleywegt, E. Birney, D. Hassabis, and S. Velankar. 2022. AlphaFold Protein Structure Database: massively expanding the structural coverage of protein-sequence space with high-accuracy models. Nucleic Acids Res. 50:D439–D444.

111. Meng, E.C., T.D. Goddard, E.F. Pettersen, G.S. Couch, Z.J. Pearson, J.H. Morris, and T.E. Ferrin. 2023. UCSF CHIMERAX : Tools for structure building and analysis. Protein Sci. 32:e4792.

112. Lomize, A.L., S.C. Todd, and I.D. Pogozheva. 2022. Spatial arrangement of proteins in planar and curved membranes by PPM 3.0. Protein Sci. 31:209–220.

113. Jo, S., T. Kim, V.G. Iyer, and W. Im. 2008. CHARMM-GUI: A web-based graphical user interface for CHARMM. J. Comput. Chem. 29:1859–1865.

114. Jorgensen, W.L., J. Chandrasekhar, J.D. Madura, R.W. Impey, and M.L. Klein. 1983. Comparison of simple potential functions for simulating liquid water. J. Chem. Phys. 79:926–935.

115. Phillips, J.C., D.J. Hardy, J.D.C. Maia, J.E. Stone, J.V. Ribeiro, R.C. Bernardi, R. Buch, G. Fiorin, J. Hénin, W. Jiang, R. McGreevy, M.C.R. Melo, B.K. Radak, R.D. Skeel, A. Singharoy, Y. Wang, B. Roux, A. Aksimentiev, Z. Luthey-Schulten, L.V. Kalé, K. Schulten, C. Chipot, and E. Tajkhorshid. 2020. Scalable molecular dynamics on CPU and GPU architectures with NAMD. J. Chem. Phys. 153:044130.

116. Huang, J., and A.D. MacKerell. 2013. CHARMM36 all-atom additive protein force field: Validation based on comparison to NMR data. J. Comput. Chem. 34:2135–2145.

117. Kroon, P., F. Grunewald, J. Barnoud, M. Van Tilburg, P. Souza, T. Wassenaar, and S. Marrink. 2024. Martinize2 and Vermouth: Unified Framework for Topology Generation. eLife, doi: 10.7554/eLife.90627.2 (preprint posted June 13, 2024).

118. Wassenaar, T.A., H.I. Ingólfsson, R.A. Böckmann, D.P. Tieleman, and S.J. Marrink. 2015. Computational Lipidomics with *insane* : A Versatile Tool for Generating Custom Membranes for Molecular Simulations. J. Chem. Theory Comput. 11:2144–2155.

119. Abraham, M.J., T. Murtola, R. Schulz, S. Páll, J.C. Smith, B. Hess, and E. Lindahl. 2015. GROMACS: High performance molecular simulations through multi-level parallelism from laptops to supercomputers. SoftwareX. 1–2:19–25.

120. De Jong, D.H., S. Baoukina, H.I. Ingólfsson, and S.J. Marrink. 2016. Martini straight: Boosting performance using a shorter cutoff and GPUs. Comput. Phys. Commun. 199:1–7.

121. Berendsen, H.J.C., J.P.M. Postma, W.F. Van Gunsteren, A. DiNola, and J.R. Haak. 1984. Molecular dynamics with coupling to an external bath. J. Chem. Phys. 81:3684–3690.

122. Kim, H., B. Fábián, and G. Hummer. 2023. Neighbor List Artifacts in Molecular Dynamics Simulations. J. Chem. Theory Comput. 19:8919–8929.

123. Wassenaar, T.A., H.I. Ingólfsson, M. Prieß, S.J. Marrink, and L.V. Schäfer. 2013. Mixing MARTINI: Electrostatic Coupling in Hybrid Atomistic–Coarse-Grained Biomolecular Simulations. J. Phys. Chem. B. 117:3516–3530.

124. Bernstein, F.C., T.F. Koetzle, G.J.B. Williams, E.F. Meyer, M.D. Brice, J.R. Rodgers, O. Kennard, T. Shimanouchi, and M. Tasumi. 1977. The protein data bank: A computer-based archival file for macromolecular structures. J. Mol. Biol. 112:535–542.

125. Stansfeld, P.J., J.E. Goose, M. Caffrey, E.P. Carpenter, J.L. Parker, S. Newstead, and M.S.P. Sansom. 2015. MemProtMD: Automated Insertion of Membrane Protein Structures into Explicit Lipid Membranes. Structure. 23:1350–1361.

126. Khelashvili, G., E. Kots, X. Cheng, M.V. Levine, and H. Weinstein. 2022. The allosteric mechanism leading to an open-groove lipid conductive state of the TMEM16F scramblase. Commun. Biol. 5:990.

127. Gumbart, J., F. Khalili-Araghi, M. Sotomayor, and B. Roux. 2012. Constant electric field simulations of the membrane potential illustrated with simple systems. Biochim. Biophys. Acta BBA - Biomembr. 1818:294–302.

128. Chen, X., N. Wang, J.-W. Liu, B. Zeng, and G.-L. Chen. 2023. TMEM63 mechanosensitive ion channels: Activation mechanisms, biological functions and human genetic disorders. Biochem. Biophys. Res. Commun. 683:149111.

129. Li, S., B. Li, L. Gao, J. Wang, and Z. Yan. 2022. Humidity response in Drosophila olfactory sensory neurons requires the mechanosensitive channel TMEM63. Nat. Commun. 13:3814.

130. Du, H., C. Ye, D. Wu, Y.-Y. Zang, L. Zhang, C. Chen, X.-Y. He, J.-J. Yang, P. Hu, Z. Xu, G. Wan, and Y.S. Shi. 2020. The Cation Channel TMEM63B Is an Osmosensor Required for Hearing. Cell Rep. 31:107596.

131. Yee, S.M., R.J. Gillams, S.E. McLain, and C.D. Lorenz. 2021. Effects of lipid heterogeneity on model human brain lipid membranes. Soft Matter. 17:126–135.

132. Chakraborty, S., M. Doktorova, T.R. Molugu, F.A. Heberle, H.L. Scott, B. Dzikovski, M. Nagao, L.-R. Stingaciu, R.F. Standaert, F.N. Barrera, J. Katsaras, G. Khelashvili, M.F. Brown, and R. Ashkar. 2020. How cholesterol stiffens unsaturated lipid membranes. Proc. Natl. Acad. Sci. 117:21896–21905.

